# Using deep learning to estimate the shifting mutational fitness landscape of SARS-CoV-2 spike along a phylogeny

**DOI:** 10.64898/2026.11.27.768997

**Authors:** Hugh K. Haddox, Jared G. Galloway, Will Dumm, Mahdi Baghbanzadeh, Jesse D. Bloom, Frederick A. Matsen

## Abstract

A longstanding challenge in evolutionary biology is to determine the fitness effects of mutations, and how they change during evolution. For SARS-CoV-2, millions of naturally evolved viral sequences are available. Bloom and Neher devised a method to estimate the effects of mutations from these sequences. The method counts how many times each mutation occurs on the branches of a phylogenetic tree and compares that count to the count expected under neutral mutation. However, the method assumes that the effect of a mutation at a site does not depend on the sequence at other sites. To estimate shifts in effects, it must divide the tree into clades chosen in advance and make a separate estimate for each clade. Here we train a deep-learning model on the entire SARS-CoV-2 tree. Because the model estimates the effects of mutations as a function of the entire sequence, it has the potential to learn a fitness landscape in which the effect of a mutation at each site depends on the sequence at all other sites. Indeed, the model learns numerous large shifts in fitness effects supported by the underlying mutation counts and in some cases independent experimental data. Further, the model’s estimated effects correlate with deep mutational scanning measurements better than those of cladespecific counts models, most notably for recent clades. Against the same measurements, our model also outperforms a recent deep-learning model that does not separate the neutral mutation rate from selection. This result suggests that separating the two helps isolate the fitness effects of mutations. We also demonstrate that, under suitable conditions, the ratio of observed to expected counts in the method of Bloom and Neher is the maximum-likelihood estimate of the selection factor in a context-free version of our model. Overall, we show how deep learning can be used to estimate a shifting fitness landscape from millions of viral sequences.

## Introduction

Six years into the SARS-CoV-2 pandemic, the virus’s spike protein continues to rapidly accumulate substitutions. The ones it has already acquired have enhanced its ability to transmit between hosts [1, 2] and have allowed it to evade host immunity and therapeutics [3–7]. A basic goal in the field is to quantify the effects of such substitutions on viral fitness [7–13] and how they have shifted over evolutionary time [14–21]. These shifts arise both from changes in external selective pressures, such as the evolving landscape of human immunity [22], and from epistasis among residues within spike [15, 17, 19]. Such knowledge is useful for interpreting past evolution and predicting future evolution for vaccine updates [9, 22–27].

To this end, an outstanding challenge is to accurately estimate the fitness effects of all possible mutations to specific SARS-CoV-2 spike sequences in a natural setting. For instance, what are the fitness effects of all mutations to the spike of XFG, a lineage that recently rose to high frequency? And how do they differ from ones to the spike of BA.2, an early Omicron variant? Deep mutational scanning (DMS) can experimentally measure the effects of individual mutations in a single genetic background on specific properties such as receptorbinding affinity or antibody neutralization [11, 12, 21, 23]. But the specific properties measured in such experiments are not the only contributors to real-world viral fitness, and the experiments are too laborious to perform in all of the different genetic backgrounds that rapidly emerge during viral evolution.

Bloom and Neher devised an approach to estimate fitness effects using a phylogenetic tree with millions of SARS-CoV-2 sequences from natural human infections [8]. Their approach counts the number of times a mutation is observed along the branches of the tree, compares that number to the counts expected under a neutral mutation process, and quantifies the mutation’s effect as the log ratio of the two.

A limitation of this approach is that it fundamentally assumes that the effects of mutations at each site are independent of the sequence composition at other sites. As a result, it cannot capture shifts in fitness effects due to epistasis or changing external selective pressures. The only way to make it capture such shifts is to partition the phylogenetic tree into clades, and then estimate fitness effects separately for each clade. But partitioning into clades is arbitrary, as evolution is continuous and fitness effects can shift along any branch. Finegrained partitions can help capture shifts, but lose statistical power due to low numbers of sequences. Coarse-grained partitions retain statistical power, but group sequences that may in fact have different fitness effects, hiding shifts within the partition.

Here, we overcome these limitations with a deep amino-acid selection model (DASM) [28], trained on a new objective in place of the likelihood of the whole child sequence. The objective is to predict which mutation occurs next on a branch [29, 41]. We show that this objective matches the counts approach of Bloom and Neher under stated conditions. A single deep-learning model is trained on the entire tree in a mutation-selection framework that allows it to learn a mutation’s fitness effect while factoring out its neutral mutation rate. The model estimates the fitness effect of a mutation as a function of the entire sequence in which it occurs. It can therefore learn an arbitrary fitness function, in which the effect of a mutation at each site depends on the sequence at all other sites, and so can capture shifts in fitness effects along the phylogeny, without the need for arbitrary clade definitions. As we show below, it also retains statistical power in sparsely sampled regions of sequence space, where clade-specific estimates lose it.

Further, our model is unique among deep-learning models of SARS-CoV-2 in that it uses a mutation-selection framework to learn fitness effects. For instance, our training objective is the same as that of antiGen [29]: both maximize the likelihood of the mutations observed along the branches of a SARS-CoV-2 tree, given the parent sequence. But the goal of antiGen is to forecast which substitutions will occur next, whereas we aim to estimate fitness effects, so our model decomposes the mutation probabilities into fitness effects and neutral mutation rates. This is important because neutral rates have been estimated to span over three orders of magnitude between mutations [30–32]. Fitness effects estimated this way correlate with deep mutational scanning measurements better than either our model’s own probabilities or those of antiGen.

## Results

### Summary of approach

Here we train a DASM on the *next-mutation objective*: given the parent sequence, the probability that a given single-nucleotide mutation is the next one to occur on a branch. This probability is the rate of that mutation divided by the summed rates of all single-nucleotide mutations of the parent, and it needs no branch length. The original DASM work [28] instead used the *child-sequence objective*: the likelihood of the whole child sequence, given the parent sequence and a jointly fitted branch length. Only the objective differs here; the architecture and the mutation–selection framework stay the same. For the purposes of this paper, “the DASM” is the DASM that we train on the next-mutation objective. The next-mutation objective parallels the counts approach of Bloom and Neher [8] and matches it under the conditions given in *Methods*. Here we focus on the spike gene, since it is the most rapidly evolving SARS-CoV-2 protein [33] and has the most experimental data available to benchmark the model.

The counts approach developed by Bloom and Neher estimates a single fitness effect for each amino-acid mutation, the same regardless of the sequence context in which the mutation occurs. The input is an UShER tree with millions of SARS-CoV-2 sequences, each of which is a consensus sequence from an infected host [34–36], with sequences at internal nodes inferred through ancestral sequence reconstruction. The approach counts the number of times a given mutation actually occurred along the tree’s branches, and compares this to the number of times it is expected to have arisen under a neutral mutation process. The mutation’s fitness effect is quantified as the log ratio of the actual to expected counts. A value of zero indicates a neutral effect, positive values indicate the mutation is beneficial (observed more than expected), and negative values indicate it is deleterious.

Our approach instead learns fitness effects that can depend on the amino acids at all other sites. This is important because at least some key mutations in spike have fitness effects that have shifted strongly during its evolution [17, 18, 20], including due to specific epistatic interactions that have been experimentally characterized [14, 16, 19, 21].

We trained a single deep-learning model on an entire UShER tree, using the approach illustrated in Figure 1A. As input, we used an UShER tree from 6 August 2026 with approximately 17 million SARS-CoV-2 sequences. We deconstructed the tree into individual branches, each of which is labeled with the spike nucleotide sequences of the parent and child nodes. We used the 1,594,774 of these branches that carried mutations in spike and passed our quality-control filters (see *Methods*), with a total of 1,708,344 mutations among them. Given the parent sequence, we trained the model to predict which mutations are present in the child sequence. As in the Bloom and Neher approach, we trained the model only on codon mutations that involve a single nucleotide change. Codon mutations involving more than one nucleotide change occur on only about 2% of the branches that carry coding mutations in spike (see *Methods*), making it difficult to estimate their effects.

**Figure 1:**
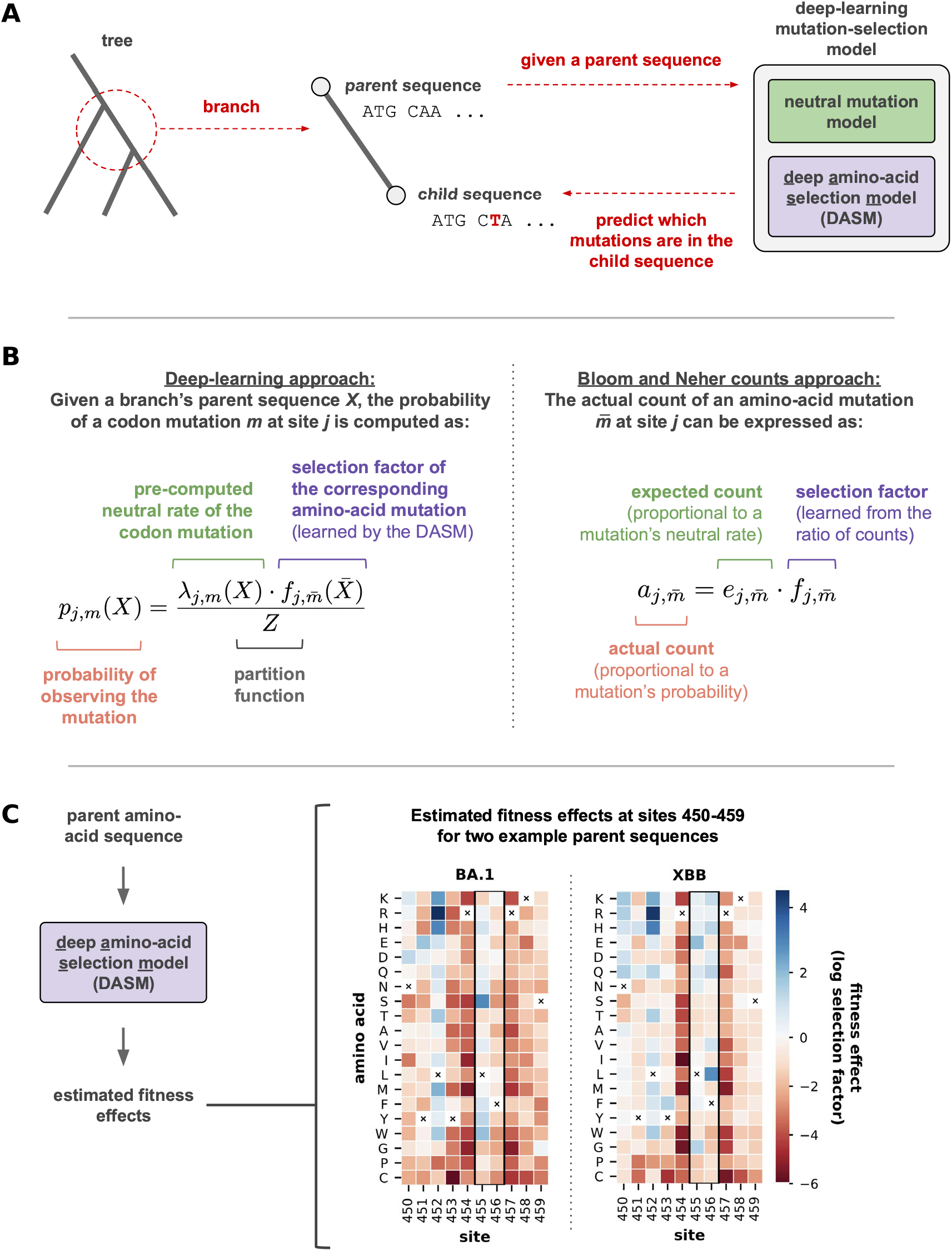
Summary of approach. **(A)** Approach to train a deep-learning mutation-selection model to predict mutations along the branches of a tree. **(B)** The left panel shows how we compute a mutation’s probability on a branch: proportional to its neutral rate multiplied by its selection factor, both conditioned on the parent sequence. The neutral rate comes from a model that we pre-trained and fixed; the selection factor comes from the DASM—a deep-learning model that adapts its estimates to the parent sequence context (*X̄* is the parent amino-acid sequence; *m̄* is the amino-acid mutation). The right panel shows the corresponding expression for the Bloom and Neher counts approach: on the branches of a tree, or of a partition of a tree, the actual count of an amino-acid mutation is its expected count under a neutral mutation process multiplied by its selection factor, which they estimate as the ratio of the two. The two expressions correspond term for term, with probabilities in place of actual counts and neutral rates in place of expected counts, but our selection factor is a function of the entire parent amino-acid sequence *X̄* , whereas theirs is context independent. **(C)** Given a branch’s parent amino-acid sequence, the DASM estimates a fitness effect (log selection factor) for each possible amino-acid mutation at 5each site in the protein. These estimates can shift between parent sequences. For example, the heat maps show estimates at sites 450–459 for the BA.1 and XBB sequences. Some estimates are substantially shifted, including at two antigenic sites (455 and 456; black rectangle). F456L, a known antibody-escape mutation, is roughly neutral in the context of the BA.1 sequence but highly beneficial in the context of the XBB sequence. An x marks the parent amino-acid identity at each site.

The probability that a mutation is observed on a branch strongly depends on both neutral mutation and selection. For SARS-CoV-2, neutral rates of nucleotide mutations have been estimated to span over three orders of magnitude depending on multiple factors, including the mutation’s type, its sequence context, base-pairing patterns in the viral RNA, and a shift in the virus’s neutral mutation spectrum upon the emergence of Omicron [30–32]. For instance, C*→*T mutations are observed at more than 100 times the rate of C*→*G mutations at synonymous sites [30]. An amino-acid mutation reached by a C*→*T change is therefore observed far more often than one reached by a C*→*G change, even when the two are equally fit, so these rates must be accounted for when inferring a mutation’s fitness effect from its counts.

Our model considers both of these factors when computing mutation probabilities. Specifically, it computes the probability of a given codon mutation at a given site as being proportional to the mutation’s neutral rate multiplied by a selection factor (Figure 1B; left formula). Neutral rates are computed using models that we pre-trained and fixed, following the neutral modeling framework of Haddox et al. [30], which models each of the above factors known to impact neutral rates. Selection factors are learned using the DASM [28]. This transformer-based deep-learning model reads the entire parent amino-acid sequence and estimates fitness effects of every amino-acid mutation at every site, including those not reachable by a single-nucleotide change from the parent codon (Figure 1C). Because the model is a transformer, it can learn to adapt its estimates to the input sequence, analogous to transformers used in other protein language models. Selection factors for synonymous mutations are fixed to a value of one, such that their probability is directly proportional to their neutral rate.

Our objective makes our model the deep-learning counterpart of the counts approach of Bloom and Neher, and each term in their expression corresponds to one in ours (Figure 1B; compare the right and left formulas). Our model’s probability of a mutation takes the place of the actual count of that mutation, and our neutral rate normalized by the partition function takes the place of the expected count. With a context-free selection factor, the maximizer of our objective is a ratio of the same form as theirs (see the last subsection of *Results* and *Methods*). Our approach differs in using a transformer to learn a fitness function where a mutation’s fitness effect depends on the parent’s sequence context. We define a mutation’s fitness effect to be its log selection factor, on the same scale as the counts-based effect above.

### The DASM learns context-dependent effects of mutations

The DASM learned large shifts in fitness effects along the UShER tree. To start, we demonstrate this using a previously characterized epistatic interaction between sites 455 and 456. A study by Jian et al. [16] found that L455F convergently arose in multiple XBB Pango lineages, but that when it did, it was almost always on the background of F456L, another mutation that arose during XBB evolution. The authors then performed experiments that helped uncover the dependence of L455F on F456L. The L455F mutation increases the resistance of XBB.1.5 spike to neutralizing antibodies in human serum, both by itself and in the presence of the F456L mutation (Figure 2A; left plot). But, by itself, L455F is highly disruptive to receptor binding. In the presence of F456L, L455F actually enhances receptor binding in addition to antibody escape (Figure 2A; right plot). So, only in the presence of F456L is L455F beneficial for both antibody escape and receptor binding.

**Figure 2:**
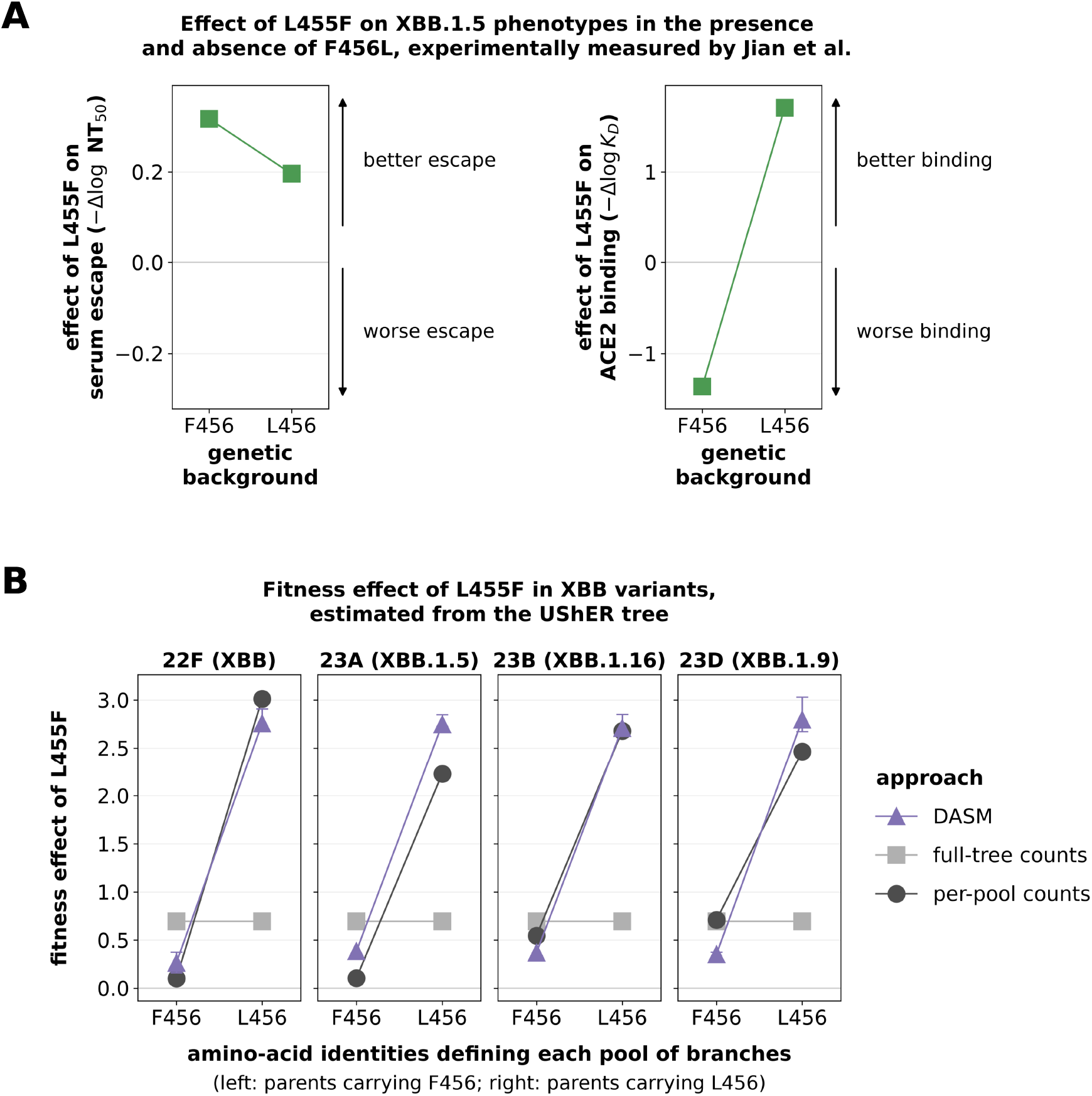
The DASM recovers an epistatic shift between sites 455 and 456. **(A)** The effect of L455F on XBB.1.5 spike phenotypes measured by Jian et al. [16]: serum neutralization (left; 50% neutralization titers from their Figure 2) and receptor binding (right; *KD* from their Figure 4). Effects are reported in two genetic backgrounds: XBB.1.5, which has an F at site 456, and XBB.1.5 with the F456L mutation. The plotted effect is the log spike phenotype without L455F minus the log phenotype with L455F, such that positive effects indicate better escape and better binding. L455F increases antibody escape in both backgrounds, but only increases binding in the presence of L at site 456. **(B)** The estimated fitness effect of L455F in four Nextstrain clades with an L at 455, on the branches whose parent carries F456 compared to the branches whose parent carries L456. Both the per-pool counts approach and the DASM approach estimate that the mutation is roughly neutral on an F456 parent and strongly beneficial on an L456 parent. The full-tree counts approach gives a single value for L455F estimated from the entire tree, and so it is completely flat across all sequence contexts. Whiskers are shown only for the DASM estimates, and span the 5th to 95th percentile of those estimates within a pool.

This epistatic shift is readily visible in the mutation counts from the UShER tree. For each of four Nextstrain clades with an L at 455, we divided the clade’s branches into two pools according to whether the parent sequence carried an F or an L at site 456, and used the counts approach to separately estimate the fitness effect of L455F within each pool (Figure 2B). Across these clades, it estimated that L455F is roughly neutral on parents carrying F456, with effects between +0.10 and +0.71, and strongly beneficial on parents carrying L456, between +2.23 and +3.01. Thus, the mutation counts capture the same striking pattern noted by Jian et al. at the level of Pango lineages: L455F occurred at a much higher rate in the presence of an L at site 456.

This is exactly the kind of context-dependent shift we would expect the DASM to be able to learn. And indeed, when we scored the parent sequences in these pools with the DASM, its estimated fitness effects showed the same shift (Figure 2B). Averaged over the parent sequences in a pool, the DASM’s effect is between +0.26 and +0.38 on the F456 pools and between +2.71 and +2.79 on the L456 pools. Fitness effects estimated from the tree are shaped by multiple phenotypes that contribute to fitness in a natural setting, including both receptor binding and antibody escape. The large positive shift in L455F’s fitness effect in the presence of F456L is consistent with L455F shifting from having a deleterious effect to a beneficial effect on receptor binding in the presence of F456L (compare Figure 2A and B), and suggests the modest decrease in L455F’s effect on antibody escape is more than counterbalanced by this large shift in its effect on binding.

The counts approach applied to the full tree cannot capture this shift, because it gives a single estimate for L455F that is the same in every sequence context (Figure 2B). We captured the shift with the counts approach only by dividing the tree’s branches into pools, split at a site we specified in advance based on prior knowledge.

The DASM learned large shifts for numerous other mutations along the tree, supported by the underlying mutation counts. To demonstrate this, we divided the tree into Nextstrain clades and fit a separate counts model to each clade, giving counts-based estimates of fitness effects for each clade, and thus of how much each mutation shifted between clades. We then used the DASM to estimate fitness effects to each clade’s founder sequence, and compared the two approaches’ estimates.

The two approaches agreed well, including for mutations that shift dramatically between clades. Figure 3A shows one such mutation, F456L—the other mutation in the epistatic pair described above—for all clades in which it had at least 10 actual or expected counts. According to the counts approach, its effect shifts from being roughly neutral in earlier clades to highly beneficial in later clades, including the XBB clades in which it reached high frequency. The DASM estimates closely track the per-clade counts estimates.

**Figure 3:**
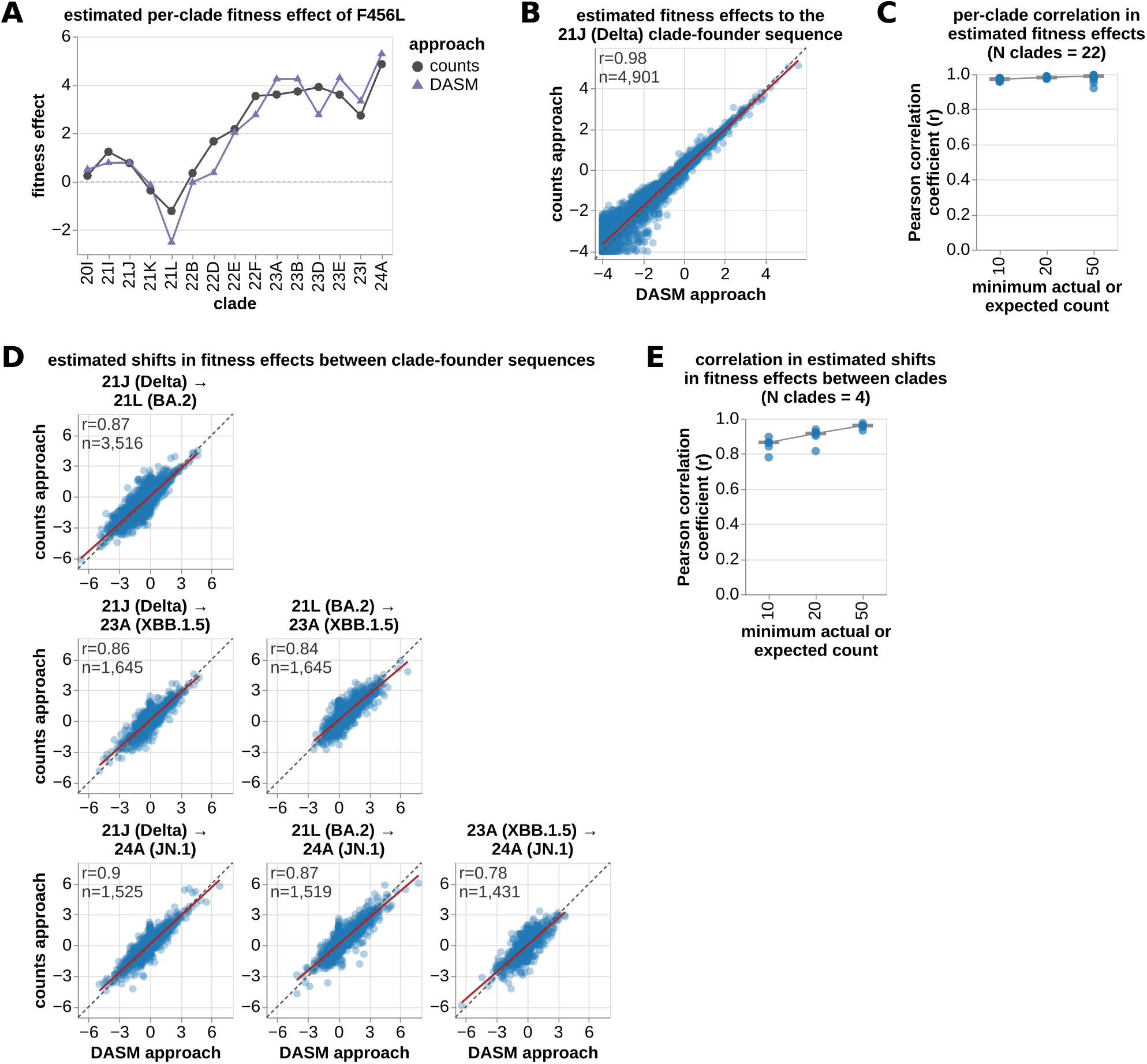
The DASM learns context-dependent effects of mutations. In all panels, counts estimates come from a separate counts model fit to each clade, and DASM estimates are fitness effects of mutations to each clade’s founder sequence, from the single DASM trained on the whole tree. **(A)** F456L is an example of a mutation that dramatically shifts between clades, according to the counts approach. The DASM predicts this shift. The plot shows clades with at least 10 actual or expected counts for F456L. **(B)** Estimated fitness effects to the Delta (21J) clade-founder sequence are highly correlated between the counts and DASM approaches. The plot shows mutations with at least 10 actual or expected counts in the clade. We clip estimated effects at a lower value of −4 because the counts approach has a limit of detection that varies between clades (see *Methods*). **(C)** Effects to clade-founder sequences are highly correlated across the 22 clades with at least 20 mutations at each counts threshold. Each dot shows the correlation coefficient for a single clade. The correlation increases as the counts threshold increases. **(D)** Estimated shifts in fitness effects for all six pairs of four clades well separated on the tree, among mutations with at least 10 actual or expected counts in both clades of a pair. A shift is the difference between a mutation’s effects in the two clades of a pair. Shifts are correlated between approaches. **(E)** As in (C), but for shifts: each dot shows the correlation coefficient for a single clade pair, and the correlation increases as the counts threshold increases.

Beyond this example mutation, the two approaches’ estimates were highly correlated within each clade, with a median Pearson *r* of 0.97 across the 22 the Pearson *r* was 0.98 among 4,901 mutations (Figure 3B; effects were clipped at values of *−*4 due to limits of detection of the counts approach; see *Methods*). Further, the median Pearson *r* increased to 0.98 and 0.99 when we progressively increased the counts threshold from 10 to 20 to 50 (Figure 3C). This increasing correlation suggests that at least some of the differences between approaches are simply due to noise. At a counts threshold of 50, the largest differences came instead from simplifying assumptions in the counts approach, including the assumption that mutational effects do not shift within clades (Table S1, Figure S1).

Because a shift is the difference of a mutation’s effects in two clades, agreement on effects should carry over to shifts, as it does for F456L. To test this, we first focused on four large clades that are broadly distributed across the tree: 21J (Delta), 21L (BA.2), 23A (XBB.1.5), and 24A (JN.1). For each pairwise combination of these clades, we estimated shifts for all mutations with at least 10 actual or expected counts in each clade. Each clade pair had a wide range of estimated shifts, and these estimates were well correlated between approaches, with a median Pearson *r* of 0.86 (Figure 3D), lower than for the per-clade effects themselves because differencing two estimates compounds their noise. As above, the median correlation increased to 0.92 and 0.96 as the counts threshold was raised from 10 to 20 to 50 (Figure 3E). The two approaches were also in strong agreement when expanding from these four clades to all pairwise combinations of the 22 clades in our analysis, as quantified by low absolute error across a range of shift values (Figure S2). The biological mechanisms underlying the observed shifts may include epistasis among residues in spike, like the interaction between sites 455 and 456 above. But they may also include changes in external selective pressures, such as human population immunity, that correlate with the presence or absence of specific mutations.

Overall, these results show that the DASM—a single model trained on the entire tree—learns shifts in fitness effects that are strongly supported by the underlying mutation counts. We emphasize that these shifts were only apparent under the counts approach after first dividing the branches into clades or pools and then separately fitting a counts model to each, whereas the DASM learns these shifts through a flexible fitness function that depends on sequence context.

### The DASM accurately estimates effects of mutations in sparsely sampled regions of sequence space

The counts approach has an inherent tradeoff, which arises because its estimates are site independent and so shifts can only be identified by dividing the tree into pieces. Fine-grained partitions are needed to resolve shifts between specific genetic backgrounds. But, the more finely the tree is divided, the more the counts approach loses statistical power in any one piece, since smaller pieces contain fewer mutation counts.

This tradeoff can severely limit the ability of counts models to accurately estimate fitness effects for individual partitions. For example, when dividing the tree into Nextstrain clades, the median clade has about 60,000 sequences (Figure 4A). This partitioning is coarse in that individual clades often combine multiple prevalent genetic backgrounds (see, for instance, Figure 2B and Figure S1). Even so, the counts within individual pieces are still limiting. In 25C (XFG), a modern clade with about 100,000 sequences, only *∼*6% of possible mutations to its founder spike sequence have at least 10 actual or expected counts when computed across that clade’s branches (Figure 4B). If we use 10 counts as a rough threshold, below which it becomes difficult for a counts model to accurately estimate a fitness effect, then a model fit to this clade would have difficulty estimating effects for 94% of possible mutations. The problem is less severe for clades with more sequences, but it does not go away. 21J (Delta) has more sequences than any other clade—over an order of magnitude more than 25C—yet only *∼*65% of mutations to its founder sequence reach this number, leaving the effects of a third of all mutations difficult to estimate accurately (Figure 4B).

**Figure 4:**
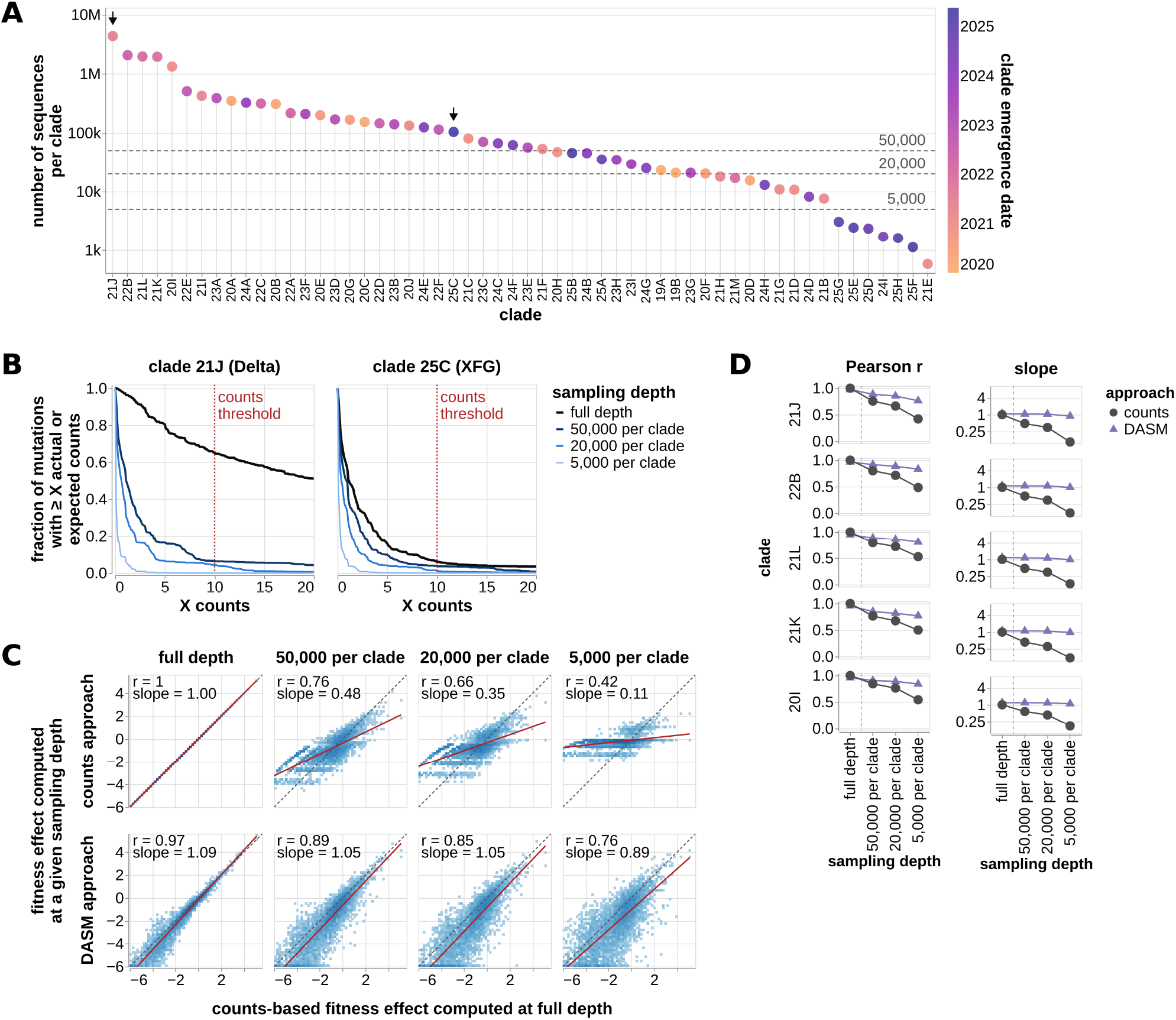
DASM estimates have higher accuracy in sparsely sampled clades. **(A)** Number of sequences per Nextstrain clade. The median clade has tens of thousands of sequences, while a few have millions. Each clade is colored by its emergence date; the dashed lines mark the three downsampling caps, and the arrows the two clades in panel B. **(B)** Fraction of mutations with at least X actual or expected counts in a given clade. In 21J (Delta), the clade with the most sequences, ∼65% of mutations have at least 10 actual or expected counts. In 25C (XFG), only ∼6% of mutations reach this number. **(C)** For the Delta clade, recovery is higher for the DASM approach at every downsampling cap. At each cap, the correlation coefficient and the slope of an OLS regression line are both closer to one. In the full-depth column the counts approach defines the ground truth, so its values are one by construction. Each plot shows only mutations with at least 10 actual or expected counts in the full Delta clade. **(D)** Recovery is higher for the DASM approach across the five clades with the most sequences, as quantified by the correlation coefficient and the slope of the OLS regression line. In panels C and D, the DASM is the 1M-parameter transformer at full depth and the 85k-parameter one at each cap, each tree being fitted with a model sized to it, and estimated effects are clipped below at −6 rather than the −4 used elsewhere, because these deeply sampled clades have a lower limit of detection.

In contrast, the DASM approach does not need to make this tradeoff. Because it uses a context-dependent fitness function, it can learn shifts without needing to divide the tree into arbitrary pieces. And because it is trained on the whole tree, its fitness function learns from all available mutation counts. We hypothesized that this would help the DASM to accurately estimate fitness effects, including in parts of sequence space where sampling is sparse and counts are limited.

To test this hypothesis, we compared the ability of the counts and DASM approaches to recover fitness effects when trained on downsampled data. For each clade, we fit a counts model to that clade alone, recorded the effects of all mutations with at least 10 actual or expected counts in the clade, and treated these as “ground truth” values. Then, we downsampled the entire tree to have a maximum of 50,000, 20,000, or 5,000 sequences per clade. This caused the number of mutation counts to substantially drop. For instance, for Delta, the caps left only a small fraction of mutations with at least 10 actual or expected counts, falling to under 1% at the smallest cap (Figure 4B). We then trained models on each tree and asked how well the model could recover the groundtruth values.

The per-clade counts models struggled. As the cap decreased, the correlation with the ground-truth values fell and the slope flattened. For Delta, the Pearson *r* fell from 0.76 to 0.66 to 0.42 across the three caps, and the slope from 0.48 to 0.35 to 0.11 (Figure 4C). These trends are expected from lower mutation counts leading to more noise. The pseudocount of 0.5 added to the actual and expected counts also tends to push estimated effects toward zero when counts are very low.

The DASM approach was much more robust to downsampling. As the cap decreased, the correlation fell by less, and the slope stayed close to one. For Delta, the Pearson *r* fell only from 0.89 to 0.85 to 0.76, while the slope stayed between 0.89 and 1.05 (Figure 4C)—even though the smallest cap leaves under 1% of Delta’s mutations with at least 10 actual or expected counts. The same pattern held across the five clades with the most sequences (Figure 4D). The DASM must therefore be learning from counts across the entire tree, and using that knowledge to inform its estimates in regions of sequence space with limited counts.

We next tested the same hypothesis against an independent benchmark: the effects of mutations measured by deep mutational scanning (DMS). We focused first on a DMS that measured the effects of mutations on spike’s ability to mediate entry of pseudoviruses into cells, in the context of the KP.3.1.1 spike sequence, which is the founder of the 24E clade [12]. Among the aminoacid mutations with fitness effects estimated by both the counts and DASM approaches—that is, those accessible by a single-nucleotide codon mutation— the DASM estimates correlate with the measured effects much better than do the estimates from a counts model fit to 24E alone, with Pearson *r* of 0.68 against 0.43 (Figure 5A). As above, this gap is expected from the low mutation counts in 24E, which leave the counts-based estimates dominated by noise and by the pseudocount.

**Figure 5:**
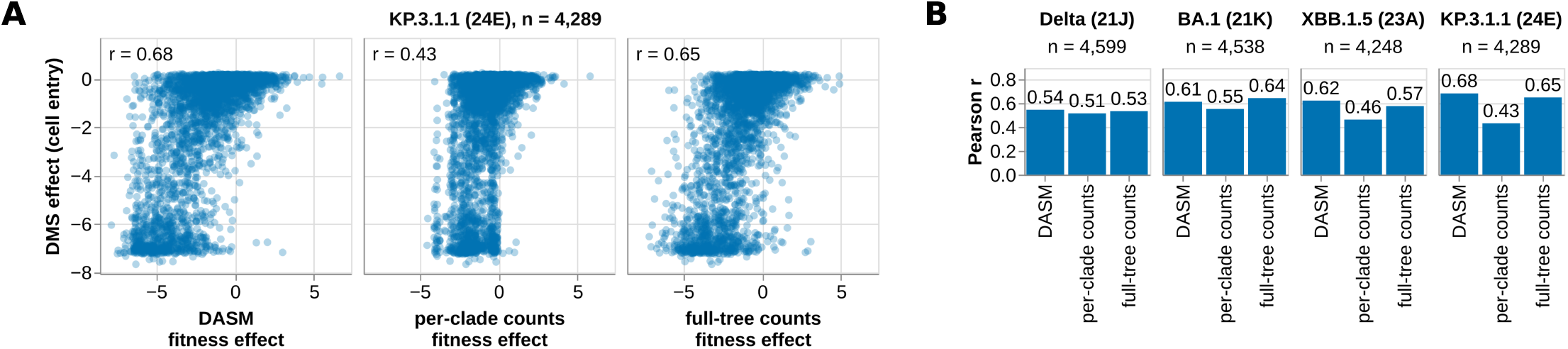
Correlation of model estimates with DMS data. **(A)** Fitness effects estimated by the DASM, by the counts approach applied to clade 24E, and by the counts approach applied to the whole tree, each against DMS measurements of spike-mediated cell entry made on the KP.3.1.1 spike. Each point is an amino-acid mutation, and Pearson *r* is given in the corner of each panel. **(B)** The same three correlations on each of the four spike backgrounds measured by DMS, ordered by decreasing number of sequences in the clade. Both panels analyze the mutations each experiment measured that the counts approach can also estimate, which are those reachable from the clade founder by a single-nucleotide codon mutation.

Consistent with this expectation, the DASM’s advantage over the per-clade counts approach shrank as the clade size increased. We saw this by evaluating the same two estimates against DMS experiments measuring the same phenotype on three older spike backgrounds, whose clades have more sequences: XBB.1.5 (clade 23A) [23], BA.1 (clade 21K), and Delta (clade 21J) [11]. The gap fell from +0.25 on KP.3.1.1 to +0.16 on XBB.1.5, +0.06 on BA.1, and +0.03 on Delta (Figure 5B). As might be expected, the per-clade counts approach does better when it has more sequences and mutations to learn from. The DASM approach does at least as well as the per-clade counts approach, but does better for more recent clades where sampling is sparser. Differences in correlation between the four experiments likely reflect factors on the experimental side, such as measurement noise, rather than factors on the model side.

The DASM’s performance on this task is roughly the same as the performance of a counts model trained on the full tree, rather than individual clades: on KP.3.1.1 the full-tree counts model reaches a Pearson *r* of 0.65 against the DASM’s 0.68, and across the four backgrounds the two never differ by more than 0.05 (Figure 5A and B). This provides additional evidence that the DASM gains statistical power by learning from all counts on the tree. We emphasize that the main advantage of the DASM over the full-tree counts model is that the DASM can detect context-dependent shifts. This advantage is unlikely to be highly visible when correlating with the DMS data, since many of the shifts along the tree are at sites of immune escape [18, 20] and thus likely shaped by immune selection, which is absent in the DMS experiments.

Overall, both analyses in this section show that counts models lose statistical power when trained on small pieces of a tree, and suggest the DASM is an effective way to overcome this limitation without sacrificing the ability of the model to learn shifts.

### Separating selection from mutation improves correlation with DMS data

Compared to other deep-learning models of SARS-CoV-2, ours is unique in that it learns fitness effects of mutations in a mutation-selection framework. Two recent preprints describe models—antiGen [29] and evoPLM-Tree [37]—that are similar to ours in that they are trained on parent-child sequence pairs taken from a SARS-CoV-2 phylogeny. antiGen shares our training objective, estimating the probability of mutations on a branch with a given parent sequence, whereas evoPLM-Tree generates a whole descendant sequence conditioned on its parent. But ours is distinct from these previous models in deconstructing its predicted probability into a pre-computed neutral rate and a learned fitness effect (Figure 1B), at least for the single-nucleotide codon mutations it is trained on.

This deconstruction is important for estimating fitness effects because probabilities can be strongly impacted by neutral rates. Below, DASM-probability means the probability that our model gives a mutation, and antiGen-probability the probability that antiGen gives it. To demonstrate this point, we consider two categories of amino-acid mutations. The first category is amino-acid mutations that are accessible by a single-nucleotide codon mutation (e.g., AAA [lysine] *→* AAT [asparagine]). The second category is amino-acid mutations that require a multi-nucleotide codon mutation (e.g., AAA [lysine] *→* CCA [proline]).

For mutations in the first category, our model estimates that two mutations with the same fitness effect can differ by over two orders of magnitude in probability, because of differences in their neutral rates (Figure 6A). Against the DMS measurements from the previous section, the fitness effects correlated better than the probabilities on all four backgrounds, with Pearson *r* higher by 0.07 to 0.13 (Figure 6B; compare DASM fitness effect with DASM-probability).

**Figure 6:**
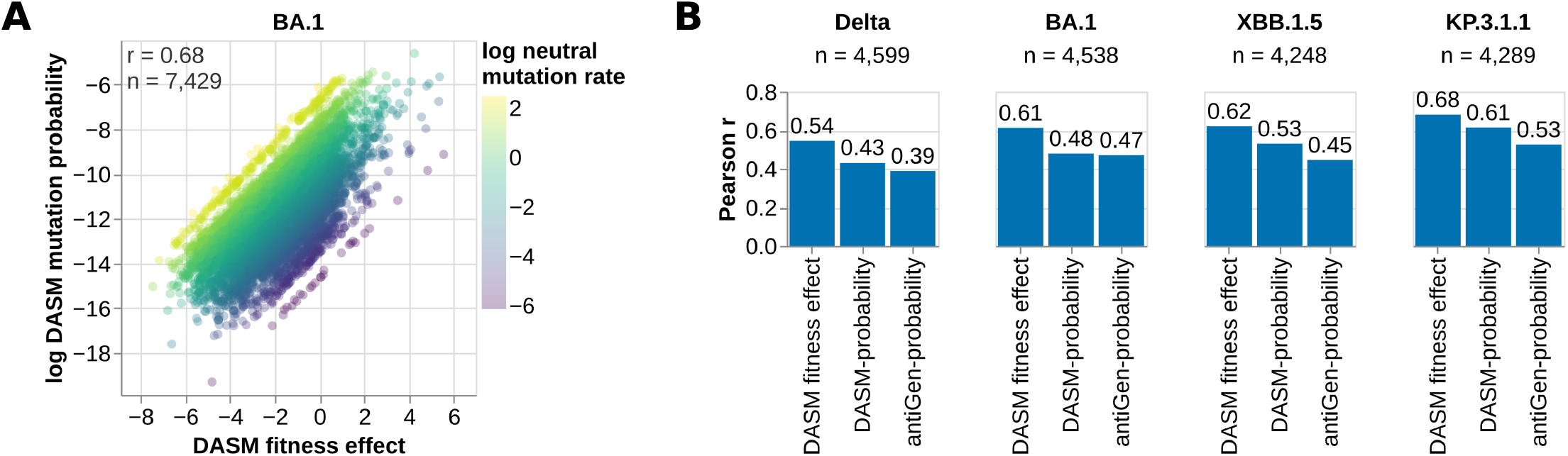
A mutation’s probability is its fitness effect displaced by its neutral rate. **(A)** For every amino-acid mutation reachable by a single-nucleotide codon mutation to the BA.1 founder spike, the fitness effect the DASM estimates against the log probability it gives that mutation on a branch out of that founder, colored by the mutation’s log neutral rate. The two quantities differ by that rate, so mutations with the same fitness effect span over two orders of magnitude in probability. Probabilities and rates are drawn as natural logarithms, so an order of magnitude is about 2.3 units on these axes. This panel draws every such mutation to the founder, not only the ones the experiment measured. **(B)** Pearson correlation with the DMS measurements on each of the four spike backgrounds, for the DASM’s fitness effect, the DASM-probability of the same mutation, and the antiGen-probability. All three are computed over the mutations the DASM gives a probability for—those reachable by a single-nucleotide codon mutation—so the populations are the same within a group.

Neutral mutation is also expected to substantially elevate the probabilities of mutations in the first category over those in the second, whose neutral rates are much lower. We were unable to search for this bias using our model, since it only computes probabilities for mutations in the first category. However, we were able to find evidence of this bias in antiGen-probabilities. antiGen, the deep-learning model referenced above, is trained to predict probabilities for all possible amino-acid mutations to a given parent sequence. To obtain an antiGen model, we trained one from the code and data that its authors made publicly available, reproducing the performance reported in their paper. For our analysis, we used the model trained on the first 5.9 years of the pandemic, the latest of the training cutoffs in that study, so that its training data contain the parent sequence of each DMS experiment, as do the DASM training data.

The KP.3.1.1 dataset shows this bias in antiGen-probabilities. We looked for it in this dataset because it measures nearly every amino-acid mutation throughout the receptor-binding domain as well as mutations elsewhere in spike; the other three experiments saturate only a small number of sites, and so measure far fewer of the mutations in the second category. The log probabilities estimated by antiGen were correlated with the DMS effects (Figure 6B). But, when we separated mutations into the two categories, we found that mutations in the first category had substantially higher probabilities than those in the second, with median log probabilities differing by 3.4 (Figure 7A; the blue dots tend to have higher probabilities than the orange dots). That difference is 1.5 times the standard deviation of the log probabilities over all measured mutations. The DMS data do not support a difference of this size: the measured effects differ between the two categories by only 0.2 standard deviations. This is the pattern expected if neutral rates are degrading the correlation between antiGen-probabilities and measured effects.

**Figure 7:**
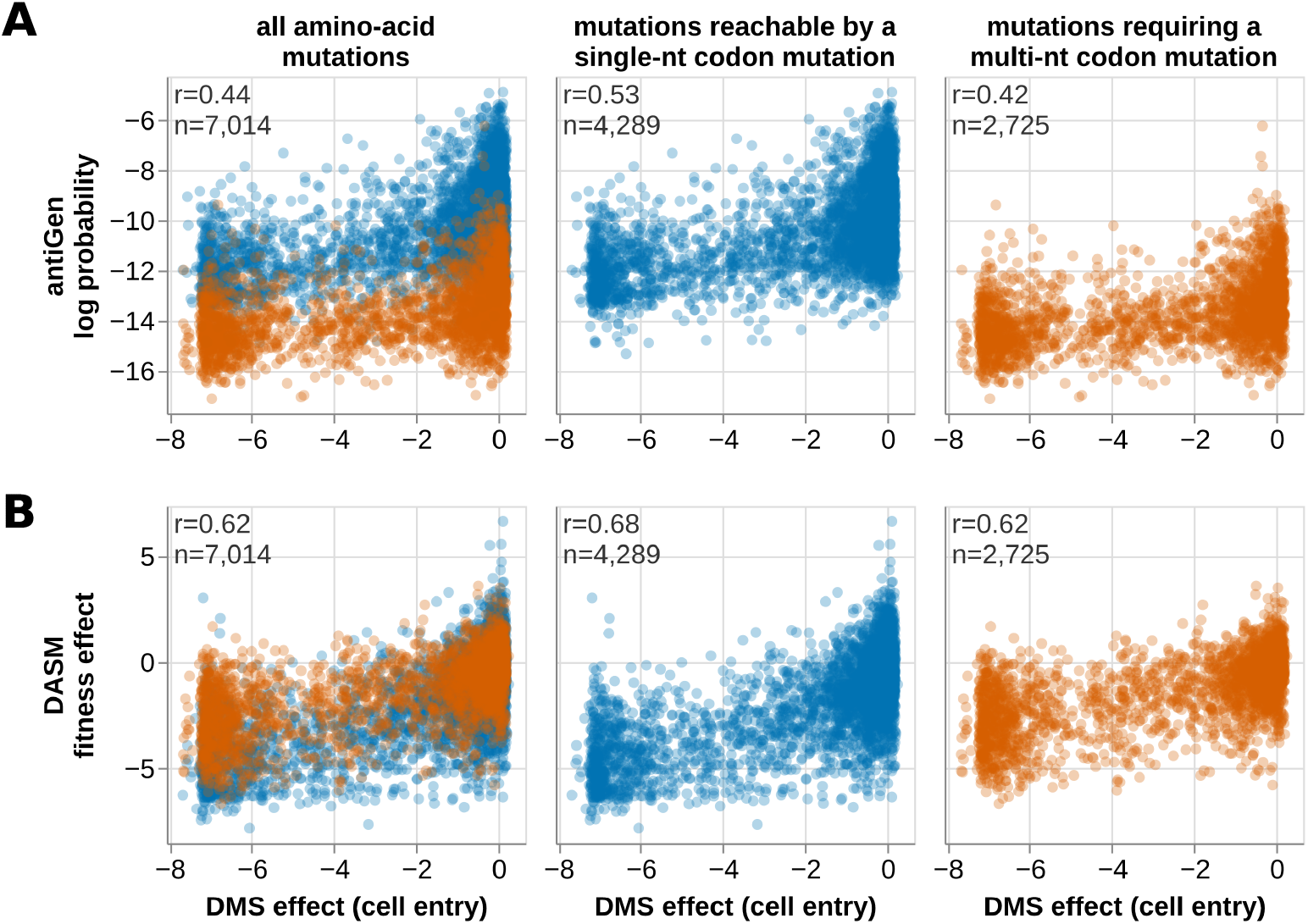
Nucleotide accessibility biases antiGen-probabilities much more than the DASM’s fitness effects. Estimates for the KP.3.1.1 founder spike against the DMS measurements of cell entry made on that background, for **(A)** the natural logarithm of the antiGen-probability and **(B)** the DASM’s fitness effect. The left column is every measured amino-acid mutation; the middle and right columns split it into the mutations reachable by a single-nucleotide codon mutation and those requiring a multi-nucleotide one. Color marks that split throughout, with the left column showing the two populations superimposed. In each cell, *r* is the Pearson correlation between the estimates and the measurements it draws, and *n* is the number of mutations.

In comparison, the fitness effects estimated with the DASM differ less between the two categories (Figure 7B). Their median difference is 0.1 standard deviations, but it runs in the opposite direction from the 0.2 standard deviations separating the DMS measurements, so the DASM departs from the DMS data by 0.3 standard deviations where antiGen departs from it by 1.4.

This result could be taken to indicate that the mutation-selection framework helps the DASM estimate the effects of mutations in the second category, but it should be taken with a grain of salt. The DASM’s estimates for these mutations do correlate with the measurements, with a Pearson *r* of 0.62 against 0.68 for the first category, even though the model is never trained on them. However, for three of the four DMS datasets, that correlation is lower than what we get by simply averaging the DASM’s estimates for the first-category mutations at the same site (Table S2). So for the second category, the transformer may not be teaching us anything beyond what it has directly learned about a site from first-category mutations.

For every DMS dataset, the DASM’s fitness effects correlate with the measurements better than antiGen-probabilities (Figure 6B, Figure 7, and Figure S3). The higher correlation could have several causes. One is the modeling framework: the DASM-probabilities agree with the measurements less well than the DASM’s fitness effects do, and the gap to antiGen narrows when we compare DASM-probabilities with antiGen-probabilities (Figure 6B). But other causes are possible, including that the DASM is trained on a larger dataset and for a substantially longer time (see *Methods*). We note that the antiGen model was not devised to estimate fitness effects in particular, and we use it here as a comparator to shed light on the difference between fitness effects and probabilities.

### The counts ratio is the maximum-likelihood estimate of a restricted case of our model

The counts approach and our model have parallel forms (Figure 1B). In the counts approach, a mutation’s actual count equals its expected count multiplied by a selection factor. In our model, a mutation’s probability on a branch is proportional to its neutral rate multiplied by a selection factor. The key difference is that our selection factor depends on the parent sequence.

In *Methods* we show that the counts ratio, before any pseudocount, is the maximum-likelihood estimate of a restricted case of our model. Intuitively, this comes down to the standard maximum-likelihood estimate of a rate as a ratio of counts. However, the statement holds only under certain conditions, which *Methods* states formally. In this restricted case, each mutation has one selection factor that does not depend on the parent sequence, and we fit that factor within one set of branches. We also evaluate every mutation at one reference sequence for the set, such as the clade founder.

In practice, our counts-based estimates deviate from this restricted case in two main ways. First, the counts-based estimates that we benchmark add a pseudocount, so the equality holds for the ratio before the pseudocount. Second, the counts approach gives a mutation the same expected count on every branch of a set, even on branches that have lost the reference residue at that site and cannot mutate there. For such sites, its expected count is too large and its estimate too low. Our model gives zero expected count to those branches. We therefore expect the counts approach to estimate the average of our model’s selection factor over the sequences in a set of branches.

For most mutations, we expect these deviations to be small, which allows us to benchmark our model against the counts approach. In our comparison, both approaches take their neutral expectation from the same pre-trained neutral models (*Methods*).

## Discussion

We devised a deep-learning method for estimating the fitness effects of mutations to SARS-CoV-2 spike and how those effects shift across the phylogeny. The method trains the DASM [28] to predict which mutation occurs next on a branch. Our model conditions on the entire gene sequence, so it learns contextdependent shifts in mutational effects that the site-independent counts approach cannot capture.

The first way our work contributes to the field is by helping to comprehensively estimate the shifting mutational fitness landscape of SARS-CoV-2 spike. The counts approach estimates this landscape one clade at a time, and only for the mutations with enough counts to support an estimate. Our approach instead makes estimates at the level of individual sequences, for every mutation. It estimated effects as well as or better than counts models fit to single clades, with the largest gain for sparsely sampled clades, perhaps because it pools counts across the whole tree. It is like a microscope that can be focused anywhere on the tree: given a sequence at a particular node, what are the effects of all possible mutations to it? A future direction would be to test how well the model does this for sequences outside its training data.

Second, we find that separating fitness effects from neutral mutation improves the prediction of deep mutational scanning measurements. A model trained to predict the next mutation has probabilities that reflect how often mutations arise as well as how selection acts on them. Previous work on influenza found that the probabilities of protein language models also reflect how easily mutations arise at the nucleotide level [38]. We investigated whether separating the two helps in two ways: by comparing with antiGen, and by calculating probabilities under the combined effect of mutation and selection in our model. antiGen trains with the same objective as our model, though over amino-acid substitutions rather than codon mutations, and without the mutation-selection decomposition. Its authors developed this objective independently of ours [29]. Fitness effects from our model correlated better with the measurements than mutation probabilities calculated in either way. A limitation is that the comparison with antiGen also differs in training data and training time, so it does not isolate the effect of the decomposition. Comparing our fitness effects with our own probabilities removes those differences, but only a model trained in amino-acid space, on the same tree and for the same number of steps, would test the decomposition directly.

Third, we show that the counts ratio, without its pseudocount, is the maximumlikelihood estimate of a restricted case of our model. This result relates the counts approach of Bloom and Neher [8] to the mutation-selection decomposition that our model takes from the DASM [28]. It holds under three conditions defined in *Methods*, and it lets us benchmark our model against the counts approach. Bloom and Neher called their log ratio a “heuristic” [8], and our result gives it a likelihood basis as an estimator of a selection factor. It also contributes to the literature that compares counting and likelihood-based estimators of selection [39, 40].

We learned of Pinion [41] in the final stages of preparing this manuscript. It uses the same mutation-selection decomposition to predict the next substitution to a parent sequence, normalized over all single-nucleotide candidates of that sequence. The two models were developed independently. Our implementation of the decomposition has been public in the netam repository since July 2025 [42], and the all-sites normalization since September 2025 [43]. Work on Pinion began in the summer of 2026 (T. Bedford, personal communication). Pinion differs from our model in four ways. It applies a Kimura fixation function to the selection factors. It fine-tunes ESMC instead of training a transformer from scratch. It co-trains with a masked language objective. It jointly trains 12 nucleotide mutation rates that are the same at every site of a parent sequence, instead of using a separately estimated neutral model with flanking effects [30]. Our approach has several limitations. Although the DASM learns a context-dependent fitness function, it cannot determine whether particular shifts in fitness effects are due to epistasis among residues in spike or due to changes in external selective pressures like human immunity that correlate with the appearance of mutations during spike’s evolution. Also, the model estimates effects through a black box, whereas the counts approach estimates them by a transparent calculation. We found good agreement between the two, but we suggest that conclusions drawn from the transformer should be validated against the underlying counts and simple baselines when possible. For instance, for aminoacid mutations reachable by a single-nucleotide codon mutation, the DASM’s estimates correlated with the DMS measurements better than counts-based estimates did; but for mutations requiring a multi-nucleotide codon mutation, which the model never sees during training, its estimates did not consistently outperform a simple average of its own estimates for the single-nucleotide mutations at that site. Like the counts approach, the model is limited by how often mutations are observed: we would not expect it to capture every shift in the tree, particularly in regions with too few sequences to show one. Its fitness effects are estimated relative to a neutral model, so they are only as accurate as that model. Epistasis between a specific pair of mutations is likewise something we would expect the model to learn only where that pair is observed often enough.

We believe that our framework may also be useful for other viruses, which are sampled far less densely than SARS-CoV-2: only hundreds of thousands of sequences are available for human influenza A virus, and far fewer for most others. Dividing such a tree into clades would leave too few counts to estimate the effects of most mutations, whereas our framework uses every sequence in the tree at once and is designed to estimate how effects shift between backgrounds. In all, we show how a deep-learning model that separates selection from neutral mutation can estimate the shifting fitness landscape of a viral protein in nature.

## Materials and methods

### Data and code availability

For all computer code we used to perform the analyses in this paper, see:

- https://github.com/matsengrp/mat-pcp: this repository processes an input UShER tree and outputs the sequence pairs used to train the DASM, and fitness effects computed using the counts approach.
- https://github.com/matsengrp/viral-dasm: this repository trains a DASM using the sequence pairs from the mat-pcp pipeline. It then performs all the analyses in our manuscript, including generation of all figures.
- https://github.com/matsengrp/s2-clade-mask: this repository builds the per-clade UShER mask from the UCSC branch-specific masking specification. The mat-pcp pipeline carries a copy of the resulting file and needs nothing else from this repository, which we provide for the mask’s provenance.

### Input data and preprocessing

Our pipeline is based on that of Bloom and Neher [8], which takes an UShER mutation-annotated tree as input and uses it to estimate fitness effects of mutations within specific clades by comparing actual mutation counts to the counts expected under neutrality. We follow it in computing the counts-based fitness effects we compare against, and in the steps that precede both analyses: subsetting the tree by clade, reading the mutations on each branch, and applying the quality control and masking described below. We additionally extract the parent and child sequences of each branch as a parent-child pair (PCP), which the DASM requires. Where they estimate expected counts from mutations at four-fold degenerate sites, we instead use the neutral mutation-rate model of Haddox et al. [30], which accounts for a mutation’s type, its sequence context, its genomic region, and whether the mutated site is paired in the genomic RNA structure. We fit the model to our input tree, and, as they did, fit one model per era: pre-Omicron and Omicron.

The specific tree that we used as input to our pipeline was a mutationannotated UShER tree with deduplicated sequences from GISAID, INSDC, COG-UK, and the China National Center for Bioinformation as of 6 August 2026 [34–36]. All GISAID sequences included in this tree and their associated metadata are available at https://doi.org/10.55876/gis8.261119vx. Accessing the sequences requires GISAID credentials, which can be obtained here: https://gisaid.org/. We are happy to share the above UShER tree upon request with anyone with GISAID access credentials. We also provide instructions for how to rerun our computational pipelines using publicly available UShER trees (see the README.md files in the root directory of each repository described above). These trees are available here: https://hgdownload.gi.ucsc.edu/goldenPath/wuhCor1/UShER_SARS-CoV-2/.

We filtered the tree’s samples and clades before analysis. The tree contains approximately 17.3 million sequences, each a consensus sequence from an infected host. We excluded samples annotated with a non-human host, samples whose collection date was missing or resolved only to the month or year, samples collected outside the window from 1 December 2019 to 6 August 2026, and samples with a genome length outside 28,000 to 32,000 nucleotides. The host annotation is absent for most samples, and we retained those. These criteria left approximately 17 million sequences. We analyzed 56 of the tree’s 58 Nextstrain clades [44], excluding 21A and 25I because of uncertainty in their founder sequences.

We extracted PCPs separately within each clade, reading the parent and child spike sequences of every branch from the tree’s ancestral sequence reconstruction and keeping the branch only when the two differed. UShER aligns all sequences to the Wuhan-Hu-1 reference when it builds the tree, discarding insertions and filling in deletions and masked sites by parsimony. Thus, the spike sequences we extracted were each 1,273 codons long, excluding the stop codon. We applied the branch-level quality control of Bloom and Neher, discarding any branch with more than four nucleotide mutations, more than one reversion to the reference base, more than one reversion to the clade-founder base, or more than one mutation within a single codon. Unlike theirs, our counts are taken over coding sites genome-wide rather than over all sites. Of the 20,972,731 branches we examined, 1,603,973 passed these filters. Among those that failed, 29,370 did so because two mutations fell within a single codon, which is 1.8% of the branches that would otherwise have passed; a branch failing several criteria is counted under the first.

We masked a small fraction of sites and mutations when training the DASM and in the analyses of our paper. These are sites that previous studies identified as highly error prone in the sequencing data [8, 45, 46], many of which are also masked when the UShER tree is built. We used the same list of sites and mutations as Haddox et al. [30], expanded to cover the clades that the tree has gained since. This amounts to 18 sites masked in every clade, and 146 further sites masked within particular clades. We also censored reversions to the reference base during training, as such mutations are frequently sequencing artifacts. At each nucleotide where a parent differs from the reference, we withheld only the mutation that restores the reference base, leaving the other mutations at that site for the model to learn from.

Finally, we removed 2,145 PCPs carrying a stop codon in the parent or child, and 7,054 PCPs whose every mutation fell at a masked position. This left 1,594,774 spike PCPs.

### A conditional probability framework for modeling mutation and selection

Here we describe a likelihood-based model inspired by Bloom and Neher [8] to infer fitness effects of mutations to a protein-coding sequence given an input phylogenetic tree. A key difference between our model and the Bloom and Neher model is that ours considers the sequence context of a mutation when estimating its effect. To do so, we first decompose the input tree into individual branches. We then train the model to predict the codon mutations on each branch given the branch’s parent sequence. For reasons described below, we only train the model to predict single-nucleotide codon mutations (i.e., codon mutations that involve one nucleotide mutation to the codon).

For a branch with a parent nucleotide sequence *X*, we define the probability of observing codon mutation *m* at position *j* to be:

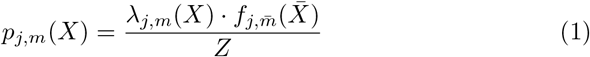

where *λ_j,m_*(*X*) is the mutation’s neutral rate given the parent sequence, *f_j,m̄_* (*X̄* ) is a selection factor for the corresponding amino-acid mutation *m̄* given the parent amino-acid sequence *X̄* , and 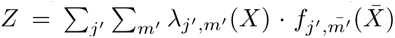 is a partition function ensuring probabilities sum to one. The inner summation term is over all possible single-nucleotide codon mutations at a given codon site in the parent sequence. The outer summation term is over all codon sites in the parent sequence. We do not consider multi-nucleotide codon mutations because they are too rare to estimate effects for, as described above; our neutral model accordingly provides rates only for single-nucleotide codon mutations. We also exclude stop codons from all consideration, excluding them from the data and from the partition function, effectively conditioning on their not appearing.

Strictly speaking, this method for computing probabilities conditions on the branch having exactly one mutation. However, we can “pretend” that this conditioning is the case for branches that have more than one mutation by giving the mutations an arbitrary order and then cutting the branches into bits that each have one mutation.

Note that our implementation does not fully realize this “cutting into bits” interpretation. A strict interpretation would update the parent sequence after each mutation and recompute the partition function *Z* for each intermediate state. Instead, our implementation uses the original parent sequence and computes a single partition function that is shared across all mutations on a branch. For a branch with *n* mutations, this means the log-likelihood contribution is 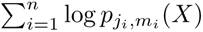 where all terms use the same *X* and *Z*. This approximation is reasonable when mutations are sparse relative to sequence length, as the partition function changes minimally between intermediate states.

The only free parameters in the model are the ones used to compute selection factors *f_j,m̄_* (*X̄* ), as we use a neutral model that we pre-trained to compute *λ_j,m_*(*X*) values, and fix the neutral model during training. Additionally, we fix selection factors of synonymous mutations to be 1, which sets a “gauge” for selection factors.

We optimize free parameters by maximizing the log-likelihood of observed single-nucleotide codon mutations:

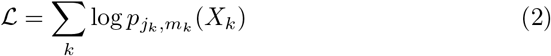

where *k* ranges over the branches of the tree, each of which has exactly one mutation *m_k_*at position *j_k_*(after cutting branches with multiple mutations into bits).

This likelihood corresponds to a forward-time model where different codon mutations compete in an exponential race. Each mutation *m* at position *j* has an instantaneous rate proportional to *λ_j,m_*(*X*) *· f_j,m̄_* (*X̄* ), and we observe whichever mutation occurs first. The fundamental result for competing exponentials is that the probability of mutation *m* winning the race is exactly its rate normalized by the sum of all rates, which gives our probability formula in Eq. 1.

Synonymous mutations play an important role in the likelihood function. Even though we fix selection factors of synonymous mutations to values of 1, the partition function will cause their *p_j,m_* values to change as the selection factors, and thus *p_j,m_* values, of nonsynonymous mutations change. Thus, selection factors of nonsynonymous mutations are fit to maximize the likelihood of both the synonymous and nonsynonymous mutations observed in the data.

The resulting selection factors can be interpreted as follows. By definition, synonymous mutations have selection factors equal to 1. We assume synonymous mutations have neutral effects. For nonsynonymous mutations, selection factors of 1 indicate the mutation has a neutral effect (which corresponds to a fitness effect of zero), selection factors *<*1 indicate the mutation is deleterious, and selection factors *>*1 indicate the mutation is beneficial.

Our model is the DASM of Matsen et al. [28], trained on a different likelihood. The DASM learns per-site selection factors with a transformer and combines them with a pre-trained neutral model. The two likelihoods differ in two main ways. First, the original likelihood covers mutations to any codon, including those requiring more than one nucleotide change, whereas ours is restricted to single-nucleotide codon mutations, since multi-nucleotide ones are too rare in these data to estimate effects for. Second, the original likelihood depends on a branch length that is optimized jointly with the model’s parameters. Because that length is fit to all the mutations on a branch, the probability the original likelihood assigns to one mutation depends on the others. Our likelihood has no branch length, and a mutation’s probability depends only on the parent sequence (Eq. 1).

### Relating the counts ratio to our likelihood

This subsection compares our likelihood with the counts approach of Bloom and Neher, and states when the counts ratio is the maximum-likelihood estimate of a restricted case of our model.

Log selection factors are conceptually similar to fitness effects inferred using the Bloom and Neher approach [8] (Figure 1B). In their approach, the selection factor of amino-acid mutation *m̄* at site *j* is inferred from the ratio of counts along the branches of a clade:

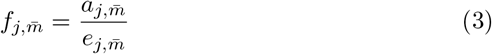

where *a_j,m̄_* is the mutation’s actual count and *e_j,m̄_* is its expected count under a neutral mutation model. As in our approach, the mutation’s fitness effect is the log selection factor, *g_j,m̄_* = log *f_j,m̄_* . Rearranging the above equation gives:

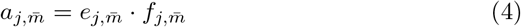

which mirrors our approach, where a mutation’s probability is proportional to its neutral rate multiplied by its selection factor (Eq. 1):

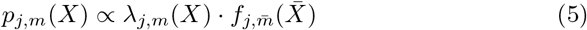

The number of times a mutation actually occurs in a set of branches (*a_j,m̄_* ) is expected to be proportional to the probability of observing it on individual branches within that set (*p_j,m_*(*X*)), assuming its fitness effect does not shift between those branches. Likewise, its expected count in the set (*e_j,m̄_* ) is proportional to its neutral rate on individual branches (*λ_j,m_*(*X*)), assuming that rate does not change between them. In both Eq. 4 and Eq. 5, the selection factor (*f_j,m̄_* and *f_j,m̄_* (*X̄* ), respectively) is a value learned from the data based on whether mutations are actually observed more or less than expected. The key difference is that our model (Eq. 5) considers the parent sequence context, while the Bloom and Neher model (Eq. 4) does not.

When selection is context free, the maximizer of our objective is a ratio of the same form as Eq. 3. When the conditions below hold and the factors are fitted within one set of branches, it equals the counts ratio before any pseudocount. To see this, drop the dependence of the selection factor on the parent sequence.

Then *f_j,m̄_* is a free scalar parameter for each site *j* and each nonsynonymous amino-acid mutation *m̄* at that site, that is, each pair of parent and mutant amino acid. The reduced model matches the mean structure of the Bloom and Neher model, with one selection factor per mutation shared by every branch in the data, whereas they fit one set of factors per clade.

Substituting Eq. 1 into Eq. 2 gives

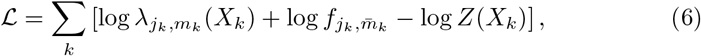

where *Z*(*X_k_*) is the partition function of Eq. 1 evaluated at the parent sequence of the branch carrying mutation *k*. Here *k* runs over mutations rather than branches. A branch with *n* mutations appears *n* times, with the same parent sequence and the same partition function each time. Only the second and third terms depend on the free parameters, so differentiating with respect to one selection factor *f_j,m̄_* gives

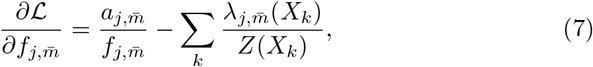

where *a_j,m̄_* is the number of mutations of that type in the data, after cutting branches with several mutations into bits. Here *λ_j,m̄_* (*X_k_*) is the total neutral rate of the codon mutations at site *j* that produce the amino-acid mutation *m̄* . Branches whose parent cannot reach *m̄* at site *j* have *λ_j,m̄_* (*X_k_*) = 0 and drop out of the sum. Branches that can reach it contribute even when the mutation did not occur there. The expected count is defined for every possible mutation, observed or not. When *a_j,m̄_* = 0, the likelihood is largest at a factor of zero, and the pseudocount keeps the log estimate finite. Setting Eq. 7 to zero gives the condition

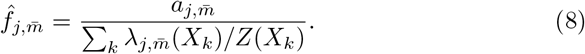

Eq. 8 has the form of Eq. 3: an actual count divided by an expected count. Equivalently, it states that the fitted model expects the observed count exactly. The denominator is the neutral rate of the mutation, summed over branches and divided on each branch by that branch’s partition function. Because each *Z*(*X_k_*) is itself evaluated at the fitted selection factors, Eq. 8 is a fixed point rather than a closed form.

Both approaches scale the expected count to the observed synonymous count, but they reach that scale differently and at a different granularity. Fixing synonymous selection factors to one, as described above, pins the overall scale, since multiplying every selection factor by a constant would otherwise leave Eq. 1 unchanged. The common scale of the nonsynonymous factors remains free, and stationarity along it gives a further identity at the optimum. The synonymous count expected under the fitted model, summed over the whole dataset, equals the observed synonymous count. Bloom and Neher calibrate separately within each clade, on four-fold degenerate sites only, with a free rate for each of the twelve mutation types rather than through the fitted factors.

Suppose that both approaches use the same neutral rates, as they do by construction in our comparison, and that the selection factors are fitted within one set of branches. The two estimators then agree exactly under three conditions. First, every mutation in the set is evaluated at one reference sequence, the founder of the clade from which the set is drawn, in place of its own parent sequence. Second, the set keeps only the mutations whose parent codon at the mutated site is the reference codon. The restricted model has no term for any other mutation, for example a C-to-T mutation at a site where an ancestor mutated A to C. We call these first two conditions together the *reference restriction*. Third, the synonymous mutations counted in the observed synonymous count are exactly the synonymous mutations in *Z*. The synonymous count sets the scale of the expected count, so a mismatch shifts every ratio in the set by one factor. Our counts pipeline and the code of Bloom and Neher both remove mutations whose parent codon differs from the reference codon. Before this removal, 1.8% of the coding mutations in our counts occur from a parent codon that differs from the clade founder’s, with a range of 0.4% to 5.8% across clades. Under the reference restriction *Z* is constant, and the identity above makes *n/Z* equal the observed synonymous count over the summed synonymous neutral rate, where *n* is the number of mutations retained in the set. That value is the scale of the counts ratio when the third condition holds. Eq. 8 then reduces to the unregularized form of Eq. 3, so the counts ratio is the maximum-likelihood estimate of a special case of our model. In our counts, the observed synonymous count exceeds the expected synonymous count by 1% to 6% in every clade. The scale uses only the synonymous mutations at the sites on which the neutral model was fit, so the third condition does not hold exactly. The two estimates therefore differ by one factor per clade.

Fitting one selection factor across several clades, each with its own reference sequence, gives a different estimate, because the synonymous identity then holds over all sets together, not within each set. In our counts, each clade has its own scale, which that clade’s synonymous counts fix without reference to the selection factors. The pooled counts-based estimate we use for the comparison with deep mutational scanning is fitted across clades in this way, so it is not exactly a maximum-likelihood estimate of our model.

With the reference restriction, the expected count of a target at site *j* does not depend on whether the parents of the branches in the set still have the reference residue there. For example, suppose two sites carry A in the founder, with equal A-to-T rates. An early A-to-T mutation at the first site spreads, so the branches below it carry T there and cannot mutate from A. The counts approach still gives both sites the same expected count, so it underestimates the selection factor at the first site. Without the reference restriction, those branches add zero (Eq. 7 and the text after it). We have not measured how large this effect is in our counts.

The log-likelihood is concave in the logarithms of the selection factors, since log *Z* is a log-sum-exp. Stationarity in *f_j,m̄_* and in log *f_j,m̄_* is equivalent for *f_j,m̄_ >* 0, so a point where Eq. 7 holds for every factor is the global maximum. The maximum is unique when every partition function has a positive synonymous rate and every nonsynonymous factor has a positive neutral rate on some branch. A factor is zero if its mutation is never observed, and the nonsynonymous factors grow without bound if the data contain no synonymous mutation.

Our model replaces the free parameter *f_j,m̄_* with the output *f_j,m̄_* (*X̄* ) of a transformer that reads the parent sequence, so the same objective gives an estimate that can differ between genetic backgrounds.

### Model architecture and training

We parameterized the selection factors *f_j,m̄_* (*X̄* ) with an amino-acid embedding concatenated with a learned positional embedding, followed by a transformer encoder and a linear layer that emits a selection factor for every amino acid at every site of the parent sequence. We trained transformers of three sizes, each with five layers, a head dimension of 8, and a dropout probability of 0.1: 84,884 parameters (4 attention heads, embedding dimension 32, feed-forward dimension 128), 292,628 parameters (8 heads, 64, 256), and 1,076,756 parameters (16 heads, 128, 512). In each, half of the embedding dimension is given to the amino acid and half to the position. Every count we give is of parameters that receive gradient during training. The DASM estimates we report are from the largest of the three transformers, except where noted. It reached the lowest validation loss of the three, although by a shrinking margin: the step from 84,884 to 292,628 parameters improved that loss by 0.008 and the step from 292,628 to 1,076,756 by 0.005, so we did not train a larger model.

We trained each model on the 1,594,774 PCPs described above, holding out a tenth of them as a validation set. For each PCP we computed, in advance, the neutral rate of every single-nucleotide codon mutation to its parent sequence, using the model fit to the era of the PCP’s clade; these rates were held fixed throughout training. We maximized the log-likelihood of Eq. 2 using AdamW in batches of 128 PCPs, with a weight decay of 10*^−^*^4^ and a learning rate of 10*^−^*^3^ for the two smaller transformers and 5 *×* 10*^−^*^4^ for the largest. We evaluated on the validation set every 2,000 steps and halved the learning rate after 10 evaluations without improvement, training until the learning rate fell below 10*^−^*^6^ and then selecting the checkpoint with the lowest validation loss. Mutations at masked sites were excluded from both the loss and the partition function.

### Counts-based fitness effects

We computed counts-based fitness effects for each clade separately, following Haddox et al. [30]. For each nucleotide mutation away from the clade founder, the actual count *a_j,m_* is the number of mutations in the clade that occur from the founder codon at that site. The expected count *e_j,m_* is the rate of that mutation under our neutral model, scaled by one factor per clade. We chose the factor so that the clade’s expected synonymous count equals its observed synonymous count at the sites on which the neutral model was fit. Those are the sites that no clade masks and at which every clade founder has the same codon and three-nucleotide context as the founder of clade 19A. This scaling absorbs how much evolution each clade has accumulated, which otherwise differs widely between them. We then summed both counts over the nucleotide mutations that encode each amino-acid mutation from the clade-founder codon, giving *a_j,m̄_* and *e_j,m̄_* , and took the fitness effect of that amino-acid mutation to be

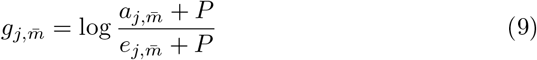

with a pseudocount *P* = 0.5, which keeps the ratio finite for a mutation that is never observed.

We also computed an estimate pooled across clades, which we used in the comparison with DMS. For each amino-acid mutation we summed *a_j,m̄_* and *e_j,m̄_* over every clade whose founder carries the same amino acid at that site, and applied Eq. 9 to those sums. Pooling the counts rather than averaging the per-clade effects weights each clade by the evidence it contributes. This is what we mean by applying the counts approach to the whole tree: the sum runs over every clade in which the mutation is defined, but not over the whole tree indiscriminately, because a clade whose founder carries a different residue at that site is not at risk for the mutation and contributes nothing.

### Estimating fitness effects within pools of branches

Both approaches can be applied to any set of branches, not only to a whole clade. We used this to estimate the effect of L455F separately on parents carrying F456 and on parents carrying L456 (Figure 2B). We took the six Nextstrain clades whose founder carries an L at site 455, and so are at risk for the mutation, and divided each clade’s branches into two pools by the parent’s amino acid at site 456. We required both pools to exist and each to reach 10 actual or expected counts, which left four clades. 23F was excluded because its founder already carries L456, so it has no F456 pool to contrast against.

Within each pool we computed the counts-based effect as in Eq. 9, refitting the scaling factor to that pool so that its expected synonymous count, summed over all synonymous mutations in spike, equals its observed synonymous count, which calibrates the expected count of each individual mutation to the amount of evolution in the pool. The expected counts of individual mutations within a pool are often far below one, and at those values the pseudocount *P* dominates the ratio and pulls the estimate toward zero, so we computed these effects without a pseudocount, applying it only where a mutation was never observed in a pool and the logarithm would otherwise be undefined. We took the DASM’s estimate within a pool to be the logarithm of the average selection factor over the distinct parent amino-acid sequences the pool contains, weighting each by the number of synonymous mutations observed on the branches it is the parent of, which measures how much of the pool’s evolutionary time that sequence accounts for. Averaging the selection factors rather than their logarithms is what makes this comparable to the counts approach, which divides a pool’s summed actual counts by its summed expected counts. The whiskers in Figure 2B span the 5th to 95th percentiles of those parent sequences’ estimates under the same weights. Alongside these per-pool estimates, Figure 2B shows the counts approach’s whole-tree estimate for L455F, pooled as described above, which is a single value and so is the same on both pools of every clade.

### Benchmarking the DASM against the counts approach

We benchmarked the DASM against the counts approach by comparing, at each clade-founder sequence, both the estimated fitness effects of individual amino-acid mutations and their shifts between clades. For the DASM we gave the model that clade’s founder amino-acid sequence as the parent sequence and took each mutation’s fitness effect to be its log selection factor; for the counts approach we took the effect computed for that clade alone (Eq. 9). Both approaches take their neutral expectation from the same pre-trained neutral models: the DASM uses them for the per-branch neutral rates *λ_j,m_*(*X*) of Eq. 1, and the counts approach uses them for the expected counts *e_j,m̄_* of Eq. 9, rescaled by one factor per clade.

We compared only those estimates the counts support, requiring a mutation’s actual or expected count in a clade to reach a threshold, and reporting thresholds of 10, 20, and 50. We excluded synonymous mutations, whose selection factor is fixed to one. We clipped both estimates below at *−*4, because the countsbased estimate has a floor that varies between clades. A mutation that is never observed takes the value log(*P/*(*e_j,m̄_* + *P* )), which is less negative the fewer counts are expected, so the limit of detection sits closer to zero in clades with fewer counts. We set the clip at *−*4, the approximate floor of the smallest clades we compare, so that a mutation below the limit of detection in two clades takes the same value in both: clipping each clade at its own floor would make a shift between clades reflect the difference between their floors rather than a change in the effect.

We report one Pearson correlation per clade, over the 22 of the tree’s 56 Nextstrain clades that retain at least 20 mutations at every threshold, so the clade set does not change between thresholds. For shifts we used four wellsampled clades spread across the tree, 21J, 21L, 23A, and 24A, giving six pairs, and defined a mutation’s shift between two clades as the difference of its effects in them. Because shifts are differences of clipped estimates, a mutation sitting at the clip in both clades has a shift of exactly zero. We then extended this comparison to every pair of those 22 clades, excluding the five pairs whose clade founders have identical spike sequences—for which the DASM’s shifts are all zero and the correlation undefined—leaving 226 of the 231 pairs. Across all pairs we report absolute error rather than correlation, because many pairs contain few mutations whose effects shift appreciably, and a correlation computed over a set of shifts that are all close to zero is dominated by noise. We pooled shifts from all pairs, binned them by the magnitude of the counts-based shift into windows of one unit, and report the median absolute difference between the two approaches’ shifts within each window, so that agreement on large shifts is not averaged together with the many small ones.

All of the estimates compared here are published with our code, in a single table: results/selection factors/dasm-1077k full depth.parquet in the viral-dasm repository, which gives both approaches’ estimates of the fitness effect of every amino-acid mutation at every spike site on every clade founder. Figure 3 is a filtering of this table.

### Detecting shifts in fitness effects within clades

We ranked the mutations of every clade by the absolute difference between the DASM’s estimate at the clade founder and the counts-based estimate for that clade, over mutations whose actual or expected count in that clade reached 50. The counts approach makes two approximations that can inflate such a difference. First, Eq. 9 adds a pseudocount *P* = 0.5 to both counts, so where the expected count is far below one the pseudocount dominates the denominator, the estimate stops tracking how small that expectation was, and a strongly beneficial mutation is understated. Second, the expected count is scaled to the whole clade while the actual count is taken only on branches whose parent still carries the founder residue, so a mutation that has already risen to high frequency had only a fraction *ϕ* of that opportunity. Table S1 reports each difference recomputed against log(*a_j,m̄_ /e_j,m̄_ ϕ*), which accounts for both. We investigated the three clades at the head of the ranking—22D, 20B, and 21K— and nine mutations within them, four of which we took from the skew away from the diagonal in Figure S1A rather than from the ranking itself.

For each clade we identified an alternative genetic background as follows. First, we found the mutation away from the clade founder that reached the highest frequency, which we call the clade’s *marker*. We measured a mutation’s frequency as the share of the clade’s total branch length on which it is present, and we set aside mutations present on over 95% of that length, which are effectively fixed and so distinguish no part of the clade from any other. Second, we found the marker’s *co-travellers*: mutations present on over half of the branch length that carries the marker. We grouped these with the marker because they often co-occur with it. The marker and its co-travellers then define the split into two pools: one set of branches carries the clade-founder amino acid at all of these sites, and the other carries the derived amino acid at all of them. In 22D the marker was R346T, present on 84% of the clade, with co-travellers F490S and K356T on 75% and 62% of the marker’s branch length. In 20B the marker was P681H, present on 17% of the clade, with co-travellers T478K and T732A on 62% and 61%. In 21K the marker was R346K, present on 55% of the clade, with no co-travellers.

We required the parent of every branch in both pools to carry the founder amino acid at the mutated site, since the mutation cannot occur otherwise, and then estimated each mutation’s effect in each pool by both approaches as described above. D574V was never observed on the 22D pool carrying R346T, K356T, and F490S, and is the one estimate here that takes the pseudocount.

### Measuring how performance degrades as sampling thins

We measured how each approach’s performance degrades as sampling thins by downsampling the tree and asking each to recover the counts-based fitness effects estimated from the full tree, which we take as ground truth throughout. We date a clade’s emergence as the earliest collection date by which 1% of its samples had been collected, rather than its first sample, so that one back-dated sequence cannot move it by months. A clade’s mutations are the amino-acid mutations reachable by a single-nucleotide change from its founder codons, excluding those to a stop, for which the DASM gives no estimate—7,504 of them for Delta’s spike. We scored each approach per clade, over the mutations whose actual or expected count within that clade reached 10 in the full tree, and over the clades holding at least 20 such mutations. We report recovery at each cap as the Pearson correlation with ground truth, together with the slope of an ordinary least-squares fit to it, since a correlation alone cannot see the compression of effects toward zero that thinning produces.

We downsampled by capping the number of sequences per clade, at caps of 50,000, 20,000, and 5,000 sequences. For a clade above the cap we drew that many of its sequences at random, without replacement, and extracted the subtree spanning them; clades at or below the cap we left untouched. These caps thin 30, 41, and 49 of the 56 clades, leaving 24%, 14%, and 3.4% of the parent-child pairs. At each cap we recomputed the counts-based effects for each clade from that clade’s own thinned subtree, and retrained the DASM from scratch on the whole thinned tree. We fitted each tree with a model sized to it: the 1,076,756-parameter transformer at full depth, and the 84,884-parameter one at each cap, which hold far less data. Little is lost by the smaller model: fitted to the full tree, it recovers the per-clade counts-based effects almost as well as the largest does, with a median Pearson *r* of 0.975 against 0.979.

### Comparing estimated fitness effects with deep mutational scanning

We compared our estimated fitness effects with deep mutational scanning measurements of spike-mediated cell entry, made on four spike backgrounds: Delta (clade 21J) and BA.1 (clade 21K), both from the same experiment [11], XBB.1.5 (clade 23A) [23], and KP.3.1.1 (clade 24E) [12]. Each experiment measures a mutation’s effect on spike-mediated entry of pseudotyped lentiviruses into ACE2-expressing 293T cells, as the log enrichment of the variants carrying the mutation between a sample taken after entry and one taken before, oriented so that higher values mean better entry and reported in Wuhan-Hu-1 site numbering. None of the four measures every amino-acid mutation to spike. Each library was designed to include the amino-acid mutations observed among natural SARS-CoV-2 sequences, which leaves out many strongly deleterious mutations, together with every amino acid at a smaller set of sites of evolutionary or antigenic interest. The KP.3.1.1 libraries also saturate the whole receptorbinding domain. We filtered each dataset as its own study filtered it. For Delta and BA.1, which come from one experiment, we kept the mutations seen in at least three variants. For XBB.1.5 we kept the mutations seen in at least three variants, measured in both replicate libraries, and with a standard error of at most 1.6. For KP.3.1.1 we kept the mutations seen in at least two variants and with a standard error of at most 2.5. We excluded mutations to a stop codon or a deletion, mutations at masked sites, and mutations at the few sites where an experiment’s wild-type residue differs from the clade founder. This left 6,143 mutations measured on Delta, 6,808 on BA.1, 6,194 on XBB.1.5, and 7,014 on KP.3.1.1.

The parent spike sequence of each experiment corresponds to the founder sequence of its clade at every unmasked position: the two carry the same amino acid wherever both carry a residue. They differ only in which positions carry a residue at all: the construct ends at residue 1252, 21 residues short of the 1,273 of full-length spike, and it carries deletions where its lineage has lost residues, while the founder carries a residue throughout, the UShER pipeline having aligned it to Wuhan-Hu-1 and filled each deletion with the most parsimonious residue. We scored the founder sequence of the matching clade with the DASM, and took the counts-based estimates to be the effect computed for that clade alone and the effect pooled across all clades (Eq. 9). We did not clip these estimates, unlike those in the comparisons above, where a common floor keeps the counts approach’s clade-varying limit of detection out of the comparison; here a floor would instead hide a difference the comparison is meant to expose.

### Scoring mutations with antiGen

The antiGen authors released their code and training data but not trained weights, so we trained the model ourselves, using commit de963f6 of https://github.com/evo-design/antiGen. Their procedure trains a series of eighteen models, each seeing only the part of the tree that predates a given date, with every subtree descending from that date held out so that predictions can be tested on it. We checked our reproduction in two ways. First, our data generation reproduces the authors’ published position-specific count matrices at each cutoff, and all of the published variant position-specific scoring matrices. This indicates that our data generation matches the authors’ exactly.

Second, across the eighteen trained cutoffs, the recall AUC and top-10% recall of our models differ from the authors’ by at most 0.0024 and 0.0116 respectively on all and variant-descended mutations, with mean differences below 0.0003 and no systematic direction. In each case, the largest difference at any cutoff is no greater than the standard deviation we measure across retrainings at the one year cutoff, using eight random seeds.

To place these differences on the scale of the quantities themselves: top-10% recall climbs from 0.40 at the earliest cutoff to 0.92 at the latest, and recall AUC from 0.80 to 0.94. Our largest difference at any cutoff is at most 0.3% of the AUC value at that cutoff and 1.7% of the top-10% recall value, and under 2% of the range each metric traverses across the cutoffs.

To compare these differences to the margins on which the paper’s comparative claims rest: on the one-year benchmark, the strongest competing baseline on both slices is the training-window PSSM, which antiGen leads by 0.036 and 0.047 recall AUC on all and variant-descended mutations respectively, while trailing by 0.095 and 0.126 on top-10% recall. The largest discrepancy between our retrained models and the authors’ is at most an eighth of the smallest of these margins.

We evaluated the model trained at the latest of those dates, 5.939 years after the tree’s root of 23 December 2019, because it is the only one whose training data contain all four DMS backgrounds. For each DMS parent sequence we supplied the corresponding sequence from the antiGen training data: the Wuhan-Hu-1 projection of the spike construct that experiment expressed, carrying the lineage’s deletions as gaps. The model returns a probability for every amino-acid substitution to that parent, and we report the logarithm of that probability. Our training of the DASM differs from antiGen’s in framework, in data, and in budget: we trained in a mutation-selection framework, on a tree with about ten times as many branches, and for 394,000 optimizer steps against the 10,000 at which antiGen’s recipe caps a run.

## Acknowledgments

We gratefully acknowledge all data contributors, i.e., the authors and their originating laboratories responsible for obtaining the specimens, and their submitting laboratories for generating the genetic sequence and metadata and sharing via the GISAID Initiative, on which this research is based (GISAID Identifier: EPI SET 261119vx, https://doi.org/10.55876/gis8.261119vx). We thank Angie Hinrichs for providing the UShER tree used in this study. We thank Trevor Bedford for useful discussions.

## Funding

This research was supported in part by grant no. R01 AI146028 from the NIH. FAM and JDB are investigators of the Howard Hughes Medical Institute. Scientific Computing Infrastructure at Fred Hutch funded by ORIP grant S10OD028685.

## Disclosures

JDB consults for Apriori Bio, Invivyd, the Vaccine Company, Pfizer, GSK, and Merck.

## Notes

https://github.com/matsengrp/viral-dasm

https://github.com/matsengrp/mat-pcp

