## Supplemental Information for "Using deep learning to estimate the shifting mutational fitness landscape of SARS-CoV-2 spike along a phylogeny"

November 27, 2026

### Tables

| Clade | Mutation | Actual count | Expected count | Fitness effect: Counts | Fitness effect: DASM | Absolute difference | Opportunity (%) | Adjusted difference |
|---|---|---|---|---|---|---|---|---|
| 22D | R346T | 145 | 0.420 | 5.06 | 8.32 | 3.25 | 15.5 | 0.61 |
| 22D | D574V | 55 | 0.872 | 3.70 | 6.94 | 3.24 | 93.1 | 2.72 |
| 22D | K356T | 56 | 0.643 | 3.90 | 7.06 | 3.16 | 43.9 | 1.77 |
| 20B | E1258D | 348 | 4.035 | 4.34 | 1.27 | 3.07 | 99.5 | 3.19 |
| 20B | D1259H | 137 | 0.775 | 4.68 | 1.64 | 3.04 | 99.9 | 3.54 |
| 21K | M177V | 463 | 27.457 | 2.81 | 0.09 | 2.72 | 99.9 | 2.73 |
| 22D | F490S | 92 | 1.850 | 3.67 | 5.84 | 2.17 | 35.2 | 0.89 |
| 24A | S31Y | 63 | 4.093 | 2.63 | 0.51 | 2.11 | 97.9 | 2.24 |
| 21K | P463T | 75 | 7.566 | 2.24 | 0.18 | 2.06 | 100.0 | 2.12 |
| 25C | V642G | 185 | 0.114 | 5.71 | 7.77 | 2.06 | 99.8 | 0.38 |

**Table S1: Mutations with the largest differences between the per-clade counts and DASM estimates of fitness effects.** For each clade we fit a counts model to that clade and evaluated the DASM on the clade-founder sequence, then took the absolute difference between the two approaches' estimates of each mutation's fitness effect. Rows are the largest of those differences, among mutations whose actual or expected count in the clade reached 50; all ten exceed 2. The actual and expected counts are $a_{j,\bar{m}}$ and $e_{j,\bar{m}}$ of Eq. 9. "Opportunity" is the percentage of the clade's branch length whose parent still carries the founder residue at that site, and "adjusted difference" recomputes the difference with that opportunity and the counts approach's pseudocount accounted for, as the *Methods* describe. Some differences are largely explained this way, as for R346T in 22D and V642G in 25C. Where one survives the adjustment, the two approaches disagree about selection, which happens when a mutation's effect shifts within the clade—something the counts approach cannot express, since it pools all of a clade's branches. Figure S1 shows the three clades at the head of this ranking, each divided into two genetic backgrounds.

| | | | Pearson $r$ with the DMS | |
|---|---|---|---|---|
| **Background** | **Category** | $n$ | **DASM** | **Site average (first category)** |
| KP.3.1.1 | First | 4289 | **0.68** | 0.61 |
| | Second | 2725 | 0.62 | **0.70** |
| | All | 7014 | 0.62 | **0.64** |
| XBB.1.5 | First | 4248 | **0.62** | 0.55 |
| | Second | 1946 | 0.45 | **0.53** |
| | All | 6194 | **0.56** | 0.54 |
| BA.1 | First | 4538 | **0.61** | 0.56 |
| | Second | 2270 | 0.42 | **0.50** |
| | All | 6808 | **0.54** | 0.53 |
| Delta | First | 4599 | **0.54** | 0.50 |
| | Second | 1544 | **0.38** | 0.35 |
| | All | 6143 | **0.51** | 0.46 |

**Table S2: The DASM's fitness effects and a per-site average of its first-category effects, each correlated with the DMS measurements.** Each row gives the Pearson correlation between one DMS experiment's measured effects on cell entry and two estimates of those effects, over the mutations the second column names. "DASM" is the model's own fitness effect for each mutation. "Site average" replaces that estimate with a single number per site: the average DASM fitness effect among the amino-acid mutations at that site that are reachable by a single-nucleotide codon mutation, which is the first category. $n$ is the number of measured mutations the row correlates. For the first category of mutations (the type the DASM saw during training), the DASM's own estimates correlate better than the site average on all four backgrounds. For the second category (the type the DASM did not see during training), the site average correlates better on every background but Delta. For all mutations, which pools the two categories, the two estimates fall within 0.02 of each other on every background but Delta, where the DASM's correlate better.

### Figures

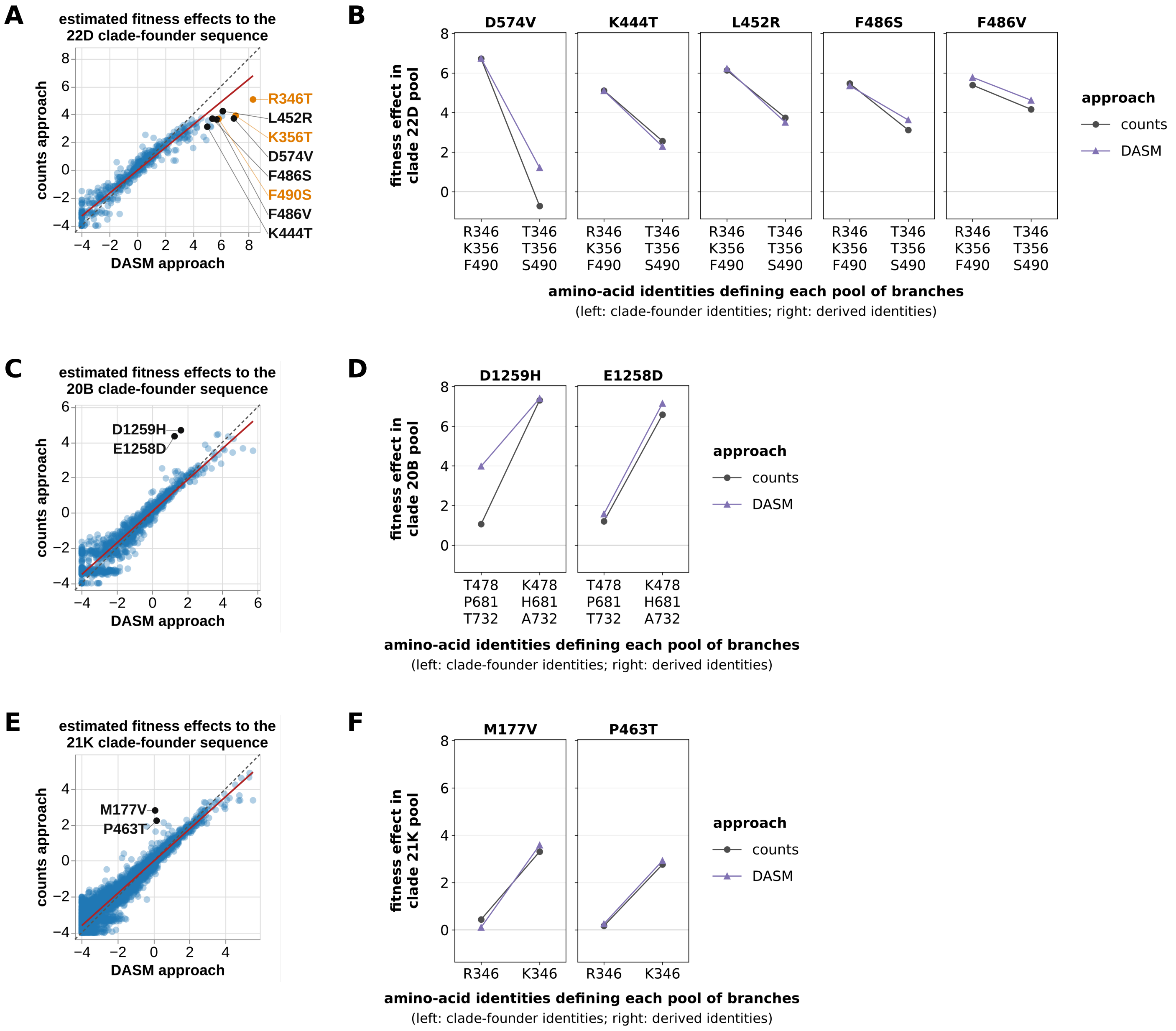


**Figure S1: The DASM identifies shifts in fitness effects within clades.** The three clades at the head of Table S1, in which the largest disagreements between the two approaches survive the adjustment described there and arise instead from shifts within the clade. **(A)** For the 22D clade, estimated fitness effects to the clade-founder sequence, counts approach against DASM approach, for mutations with at least 10 actual or expected counts in the clade. Black labels the mutations followed up in B. These include mutations from the ranking in Table S1, together with four others in 22D that show the same skew away from the diagonal but are not in that table: F486S and L452R differ between the two approaches by less than any of the ten rows listed there, and K444T and F486V do not reach its count threshold of 50. Orange labels the three mutations that define the two backgrounds in B, which rose to high frequency in this clade; the counts approach understates these because they had only a fraction of the clade's opportunity, not because their effects shift. **(B)** Each black-labeled mutation's effect in each of the two backgrounds, by both approaches, with the backgrounds named by their parent amino-acid identities at the sites that define them. Counts-based estimates in B, D, and F omit the pseudocount of Eq. 9, which improves the agreement between the two approaches; it is applied only where a mutation is never observed on a background, as D574V is on the background carrying all three mutations. **(C, D)** The same two panels for the 20B clade, and **(E, F)** for the 21K clade.

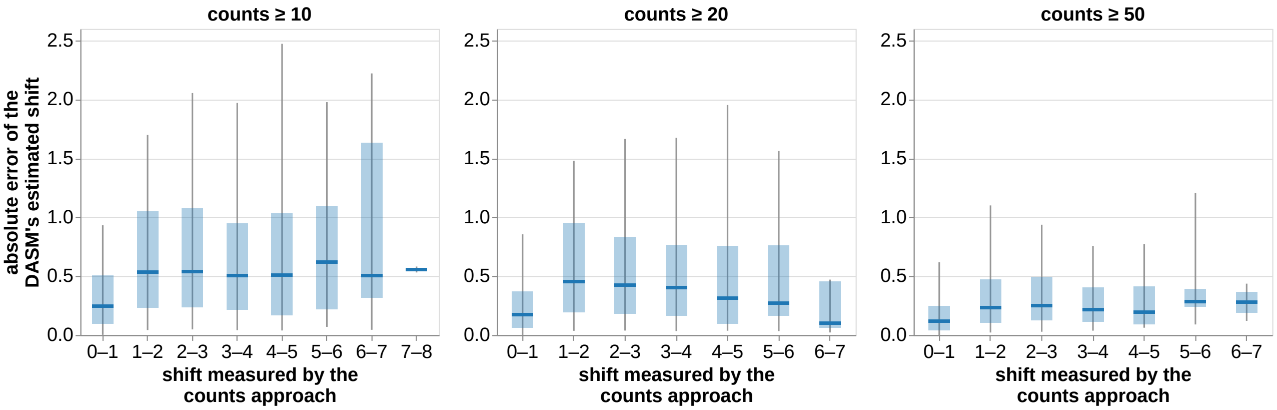


**Figure S2: Error in the DASM's estimated shifts, across all pairs of 22 clades.** For every pair of the 22 clades and every mutation with at least the stated number of actual or expected counts in each clade, we estimated the shift in the mutation's fitness effect between the two clades by both approaches, and took the absolute difference between the two estimates. Mutations are grouped into windows of one nat by the magnitude of their counts-based shift, which the x axis labels. Each box spans the 25th to the 75th percentile of absolute error within its window, the horizontal line is the median, and the whiskers span the 5th to the 95th percentile. The three panels raise the counts threshold from 10 to 20 to 50, as Fig. 3E does. At a threshold of 10, the comparison covers 176,120 (mutation, clade pair) combinations drawn from 226 of the 231 pairs, and the median absolute error stays between 0.25 and 0.62 across the windows, although the spread widens with the size of the shift. We quantified this agreement by absolute error rather than by Pearson $r$ because many clade pairs have few mutations with large shifts, so a correlation computed within such a pair would be taken over a narrow range of shifts.

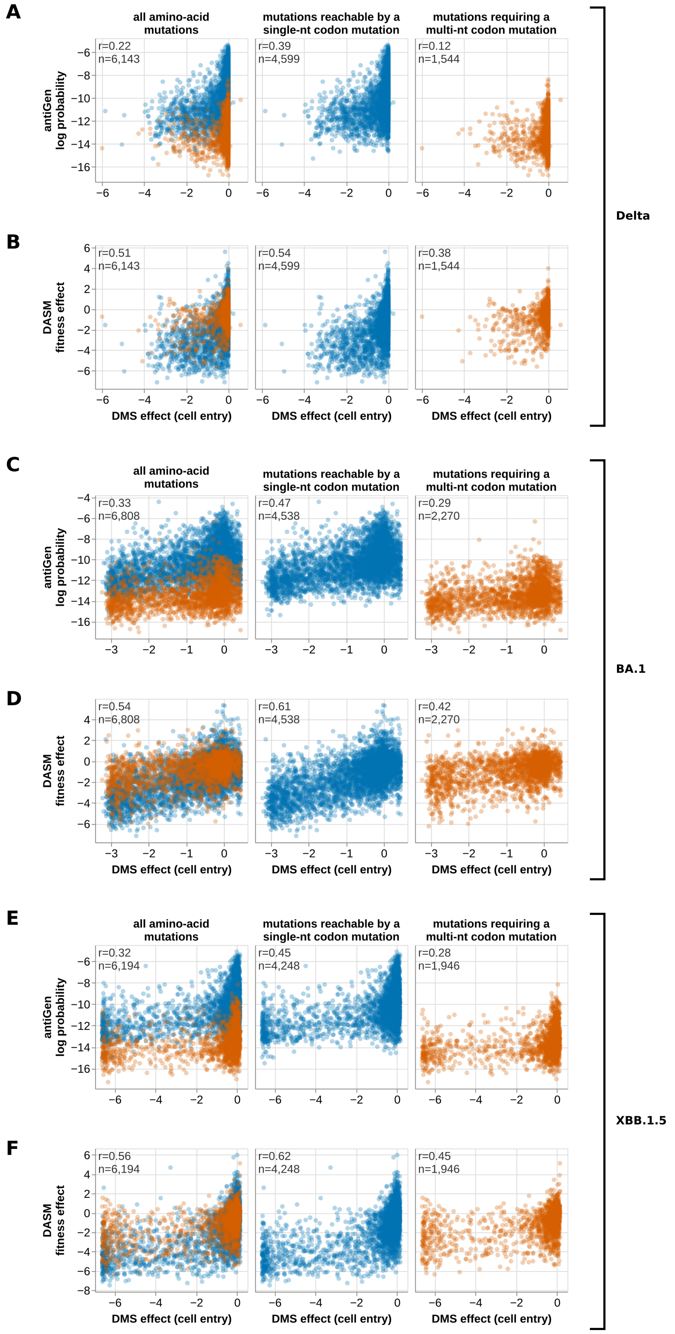


**Figure S3: Nucleotide accessibility biases antiGen-probabilities more than the DASM's fitness effects, on the other three DMS backgrounds.** As Fig. 7, for the three experiments that figure does not show, bracketed by the background each was measured on: **(A, B)** Delta, **(C, D)** BA.1, and **(E, F)** XBB.1.5. Within each pair, the upper panel is the natural logarithm of the antiGen-probability and the lower is the DASM's fitness effect. In each cell, $r$ is the Pearson correlation between the estimates and the measurements it draws, and $n$ is the number of mutations. On each background, antiGen's median log probability is higher for the first category of mutations than for the second by 1.6, 1.5, and 1.6 standard deviations of the log probabilities over all measured mutations, whereas the DMS measurements separate the two categories by 0.1 standard deviations or less. The DASM's fitness effects differ between the categories by at most 0.3 standard deviations, and in the opposite direction from the measurements, so the DASM departs from the DMS data by 0.3, 0.4, and 0.1 standard deviations where antiGen departs from it by 1.6, 1.4, and 1.6.

### References
